# Coming together – symbiont acquisition and early development of *Bathymodiolus* mussels

**DOI:** 10.1101/2020.10.09.333211

**Authors:** Maximilian Franke, Benedikt Geier, Jörg U. Hammel, Nicole Dubilier, Nikolaus Leisch

## Abstract

Symbiotic associations between animals and microorganisms are widespread and have a profound impact on the ecology, behaviour, physiology, and evolution of the host. Research on deep-sea mussels of the genus *Bathymodiolus* has revealed how chemosynthetic symbionts sustain their host with energy, allowing them to survive in the nutrient-poor environment of the deep ocean. However, to date, we know little about the initial symbiont colonization and how this is integrated into the early development of these mussels. Here we analysed the early developmental life stages of *B. azoricus, “B”. childressi* and *B. puteoserpentis* and the changes that occur once the mussels are colonized by symbionts. We combined synchrotron-radiation based µCT, correlative light and electron microscopy and fluorescence in situ hybridization to show that the symbiont colonization started when the animal settled on the sea floor and began its metamorphosis into an adult animal. Furthermore, we observed aposymbiotic life stages with a fully developed digestive system which was streamlined after symbiont acquisition. This suggests that bathymodiolin mussels change their nutritional strategy from initial filter-feeding to relying on the energy provided by their symbionts. After ∼35 years of research on bathymodiolin mussels, we are beginning to answer fundamental ecological questions concerning their life cycle and the establishment of symbiosis.

## Introduction

Mutualistic interactions between hosts and their microbiota play a fundamental role in the ecology and evolution of all animal phyla. By associating with microorganisms, animals can expand their metabolic capabilities and colonize habitats they could not live in on their own [1]. A prime example for such symbioses are bathymodiolin mussels, which occur worldwide at cold seeps and hot vents on the ocean floor. Mussels of the genus *Bathymodiolus* house chemosynthetic bacteria in their gills, in cells called bacteriocytes [2]. In these nutritional symbioses, the bacteria use reduced compounds in the vent and seep fluids, to gain energy for the fixation of carbon and provide nutrition to their hosts. Two types of symbionts dominate *Bathymodiolus* mussels, sulphur-oxidizing (SOX) symbionts, whose main source of energy are reduced sulphur compounds, and methane-oxidizing symbionts (MOX), which gain their energy from oxidizing methane [3].

The transmission of symbionts from one generation to the next plays a central role in the ecology and evolution of mutualistic associations. Symbionts can be transmitted vertically from parent to offspring, intimately tying them to the reproduction and development of their host. Alternatively, in horizontal transmission, symbionts are recruited each generation anew from the environment, entailing that the symbiotic partners are physically separate from each other before the symbiosis is established. In many hosts that rely on horizontal transmission, the acquisition of symbionts induces drastic morphological and developmental changes [4]. These can range from tissue rearrangement, as in the squid-vibrio symbiosis [5], to the development of an entirely new bacteria-housing organ, like the trophosome in the hydrothermal vent tubeworm *Riftia pachyptila* [6].

Although *Bathymodiolus* mussels have been studied for over 35 years, very little is known about how their symbionts are transmitted, at which developmental stage the symbionts colonize the mussels, and how symbiont acquisition affects the development and body plan of the mussels [7]. It is assumed that the symbionts are transmitted horizontally, based on phylogenetic studies that showed a lack of cospeciation between hosts and symbionts, and morphological studies in which symbionts were not observed in the mussels’ reproductive tissues [8-11]. In closely related “*Idas*” mussels, light and fluorescence microscopy analyses showed that these remained aposymbiotic until the juvenile stage, indicating horizontal transmission in these species [12, 13]. However, definitive evidence of an aposymbiotic early life stage of *Bathymodiolus* mussels is lacking. The earliest life stages found so far were in the late larval phase, and these were already colonized by symbionts, with a well-developed symbiotic habitus that was indistinguishable from adult mussels [14]. Given that early life stages of aposymbiotic *Bathymodiolus* have not yet been described, fundamental questions in the acquisition of symbionts in these hosts have remained unanswered, including at which developmental stage the mussels acquire their symbionts, if the SOX and MOX symbionts colonize their hosts at the same time, and which developmental changes occur in the mussels at the onset of symbiont colonization.

In this study, we were fortuitous in discovering very early, aposymbiotic life stages of three *Bathymodiolus* species, two from hydrothermal vents on the Mid-Atlantic Ridge (MAR), *B. puteoserpentis* and *B. azoricus*, and one from cold seeps in the Gulf of Mexico, “*B”. childressi* (also referred to as *Gigantidas childressi* e.g. [15]). We used a correlative imaging approach by combining synchrotron-radiation based micro-computed tomography (SRμCT), correlative light (LM) and transmission electron microscopy (TEM), and fluorescence in situ hybridization (FISH) to analyse the early life stages of *Bathymodiolus* mussels and compare them to their shallow-water relatives, the blue mussel *Mytilus edulis* [16]. This approach allowed an integrative analysis from the level of the whole animals down to single host and symbiont cells of the processes involved in symbiont colonization and its effects on the host body plan [17, 18].

## Methods

### Sampling and fixation

All deep-sea mussels were collected with remotely operated vehicles:

*B. puteoserpentis* from the Semenov vent field on the MAR at 2447 m depth with the German Research Vessel (RV) Meteor during cruise M126 in 2016, *B. azoricus* from the Bubbylon vent on the MAR at 1002 m depth during the RV Meteor cruise M82-3 in 2010, and *“B”. childressi* at the Mississippi Canyon site 853 in the Gulf of Mexico at 1071 m depth with the RV Nautilus (Ocean Exploration Trust) during cruise NA58 in 2015 (detailed information in Table S1). *M. edulis* were collected in the Baltic Sea at a site close to Kiel, Germany (54.394 N, 10.190 E) at 1.5 m water depth. Upon recovery, specimens were fixed for morphological analysis and FISH in 2% paraformaldehyde (PFA) in phosphate buffer saline (PBS), and for morphological analysis and TEM in 2.5% glutaraldehyde (GA) in PHEM buffer (piperazine-N, N′-bis, 4-(2-hydroxyethyl)-1-piperazineethanesulfonic acid, ethylene glycol-bis(β-aminoethyl ether and MgCl_2_) [19]. After fixation, samples were stored in the corresponding buffers (PFA: ethanol/PBS; GA: PHEM).

### Shell measurements

Mussels were photographed with a stereomicroscope (Nikon SMZ 25, Nikon Japan) equipped with a colour camera (Nikon Ds-Ri2, Nikon Japan) and the software NIS-Elements AR (Nikon Japan). Measurements were recorded as shown in Figure S1c (Table S2) and shell margin limits were identified by their unique coloration (Figure S1d).

### Histological analysis

PFA-fixed samples were decalcified with ethylenediaminetetraacetic acid (EDTA). For histological analysis, samples were post-fixed with 1% osmium tetroxide (OsO_4_) for 1 h and embedded in low-viscosity resin (Agar Scientific, UK). Sections of 1.5-µm thickness were cut with a microtome (Ultracut UC7 Leica Microsystem, Austria) and stained with 0.5% toluidine blue and 0.5% sodium tetraborate. For TEM, semi-thin sections were mounted on a resin block, ultra-thin (70 nm) sections were cut on a microtome (Ultracut UC7 Leica Microsystem, Austria) and mounted on formvar-coated slot grids (Agar Scientific, United Kingdom) [18]. Sections were contrasted with 0.5% aqueous uranyl acetate (Science Services, Germany) for 20 min and with 2% Reynold’s lead citrate for 6 min. For details see Supplementary Methods.

### Fluorescence in situ hybridization

PFA-fixed *B. puteoserpentis* were dehydrated, embedded in paraffin and sectioned at 5–10 μm thickness on a Leica microtome RM2255 (Leica, Germany). Sections were de-waxed, rehydrated, and air-dried. FISH was performed using double-labelled specific probes targeting the 16S RNA of the SOX and MOX symbionts as well as a general probe targeting all bacteria (Table S3) [20]. For details see Supplementary Methods.

### Microscopy

Light microscopy (LM) analyses were performed using a Zeiss Axioplan 2 (Zeiss, Germany) equipped with an automated stage, two cameras (Axio CAM MRm Zeiss and Axio CAM MRc5 Zeiss, Germany) and an Olympus BX61VS (Olympus, Tokyo, Japan) slide-scanner equipped with an automated stage and two cameras (Olympus XM10, Tokyo, Japan and Pike F-505C, Allied Vision Technologies GmbH, Stadtroda, Germany).

Fluorescence microscopy analyses were performed using an Olympus BX53 compound microscope (Olympus, Tokyo, Japan) equipped with an ORCA Flash 4.0 camera (Hamamatsu Photonics K.K, Hamamatsu, Japan) using a 40x semi-Apochromat and a 100x super-Apochromat oil-immersion objective and the software cellSens (Olympus, Tokyo, Japan), and a Zeiss LSM 780 confocal laser-scanning microscope (Carl Zeiss, Jena, Germany) equipped with an Airyscan detector (Carl Zeiss, Jena, Germany) using a 100× Plan-Apochromat oil-immersion objective and the Zen-Black software (Carl Zeiss, Jena, Germany). For details see Supplementary Methods.

Ultra-thin sections were imaged at 20–30 kV with a Quanta FEG 250 scanning electron microscope (FEI Company, USA) equipped with a STEM detector using the xT microscope control software ver. 6.2.6.3123.

### SRμCT measurements

SRμCT datasets were recorded at the Deutsches Elektronen-Synchrotron (DESY) using the P05 beamline of PETRA III, operated by the Helmholtz-Zentrum Hereon (Geesthacht, Germany [21]). The x-ray microtomography setup at 15–30 keV and 5× to 40× magnification was used to scan resin-embedded, OsO_4_-contrasted samples with attenuation contrast and uncontrasted samples in PBS-filled capillaries [22], with propagation-based phase contrast. Scan parameters are summarized in Table S4. The tomography data were processed in Matlab. Custom scripts implementing a TIE phase-retrieval algorithm and a filtered back projection, implemented in the ASTRA toolbox, were used for the tomographic reconstruction [22-25]. SRµCT models were used to ground truth the volume calculations from histological section series by measuring sectioning-induced tissue compression, which was on average 7%.

### Image processing and 3D visualization

Histograms and white balance of microscopy images were adjusted using Fiji ver. V1.52p and Adobe Photoshop CS5 and figure panels were composed using Adobe Illustrator CS5 (Adobe Systems Software Ireland Ltd.). LM-images were stitched and aligned with TrackEM2 [26] in Fiji ver. V1.52p.

Threshold-based and manual segmentation were used to generate 3D models from LM and µCT datasets from representative individuals in Amira 6.7.0 (ThermoFisher Scientific). Co-registration between µCT and LM datasets and between LM and TEM datasets was carried out in Amira [18]. For details see Supplementary Methods.

## Results

We analysed developmental stages of three species of *Bathymodiolus* mussels from aposymbiotic pediveligers to symbiotic adults, to determine at which stage the symbionts colonize their hosts and reveal the developmental modifications that these mussels have evolved to adapt to their symbiotic lifestyle (Figure 1). Because the names for larval stages of bivalves have not always been used consistently, we here define the stages we analysed. The earliest life stages in our study were at the last planktonic larval stage -the pediveliger. Once settled on the seafloor, the animal initiates its metamorphosis from a planktonic to a benthic lifestyle and enters the plantigrade stage of development. While metamorphosing, the plantigrade degrades the velum, the larval feeding and swimming organ, and develops into a post-larva. During the post larval stage, the animal secretes the adult shell and once the ventral groove of the gills is formed [27], it enters the juvenile stage. Once the gonads have developed, the mussel becomes an adult [16].

**Figure 1.**
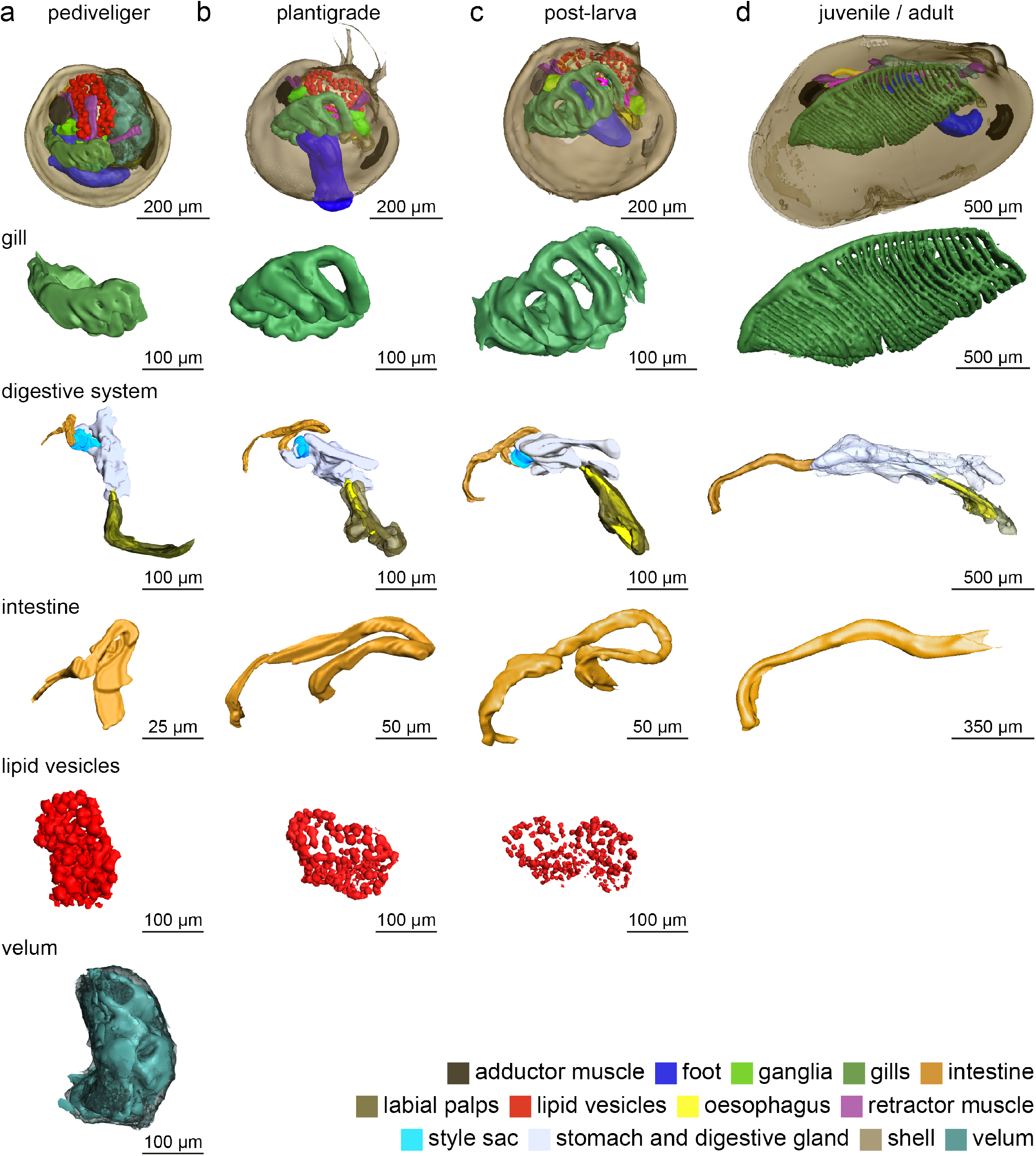
Three-dimensional visualization of *B. puteoserpentis* developmental stages analysed in this study. Note the different scale bars in a–c and d. The three-dimensional data analysed was obtained by threshold-based and manual segmentation of histological section series and SRµCT measurements.

### Identification of developmental stages

We analysed 259 *Bathymodiolus* individuals from three species that ranged from 370 µm to 4556 µm shell length (129 *B. puteoserpentis*, 124 *B. azoricus* and 6 “*B”. childressi;* Table S2, Table S5). The earliest developmental stages we recovered were pediveligers, with shell lengths of 366 µm to 465 µm. However, shell dimensions alone were not sufficient for distinguishing between developmental stages, which could only be identified through detailed analyses of the mussels’ soft body anatomy (Figure S1a and b). These morphological analyses revealed that the shell sizes of 58 pediveligers and plantigrades overlapped with those of the smallest post-larva stages (Figure S1f).

### Morphological characterization of *Bathymodiolus* developmental stages

Among the samples we collected, *B. puteoserpentis* specimens were best preserved and covered the widest range of developmental stages. We therefore focussed our detailed morphological analyses on *B. puteoserpentis*, and compared these findings with selected *B. azoricus* and *“B”. childressi* stages (Figure S2 and S3). In the following, we describe the shared morphological features of all three species, unless specified otherwise.

The pediveliger larvae were characterized by the presence of a velum (the larval swimming and feeding organ), a fully developed digestive system, a foot with two pairs of retractor muscles and two gill baskets (Table S6, Figure 1a and Video S1, Figure S4 a and d). The digestive system consisted of the mouth, oral labial palp, oesophagus, stomach, two digestive glands, gastric shield, the style sac with crystalline style, mid gut, s-shaped looped intestine, and an anal papillae (Figure 1a and Figure S5). The epithelial cells of the stomach contained membrane-bound lipid vesicles with an average diameter of 15.8 µm (75 lipid vesicles measured in 3 mussels; Figure 1 and Figure S5). The gill baskets on each side of the foot (Figure 1a and Figure S5) consisted of three to four single gill filaments in *B. puteoserpentis* and five in *B. azoricus* and *“B”. childressi* (Figure S2 b and c). These filaments form the descending lamella of the inner demibranch in later life stages (Figure S5, Video S1). For further details, see Supplementary Note 1.

In plantigrades, the developmental stage after the pediveliger, the first steps of metamorphosis from a planktonic to a benthic lifestyle were visible in the degradation of the velum (Figure S4) and the appearance of byssus threads (Figure S6). In this stage, rearrangements of all organs occurred, for example, the alignment of the growth axis of the gill ‘basket’ with the length axis of the mussel (Figure 1a-c, Figure S7 and Video S2). The number of gill filaments increased by one in all species (*B. puteoserpentis*, 5; *B. azoricus*, 6; and *“B”. childressi*, 6; Figure S3). Furthermore, gill filaments separated from each other, increasing the gaps between them from 47 µm to 120 µm (Figure 1c). These changes in gill morphology indicate a functional shift from a purely respiratory organ to a filter feeding and respiratory organ. In the digestive tract, the number and volume of lipid vesicles decreased from 12.8% of the soft-body volume in pediveligers to 3.9% in plantigrades, 1.5% in post-larvae (mean diameter 8.37 µm, *n* = 60), and were no longer present in juveniles and adults (Figure 1a-d and Table S6). The post-larval stage began once the mussels secreted the dissoconch and completed metamorphosis (Figure S1 and Figure S8). As the mussels transitioned from the post-larval to the juvenile stage, the digestive system straightened (Figure 1c-d, Figure S9 and Video S3 and S4). This morphological change was most prominent in the intestine, which went from a looped to a straight shape and remained straight in all later developmental stages (Figure 1c-d and Figure S9). For further details see Supplementary Note 2.

### Establishment of the symbiosis

Central to an accurate assessment of symbiont colonization and symbiont-mediated morphological changes was our combined approach of correlative SRµCT, FISH, and light and electron microscopy (Video S5, Figure S10). This allowed us to rapidly screen whole animals yet achieve the resolution needed to identify the colonization of single eukaryotic cells by symbiotic bacteria. We first searched for a morphological characteristic that was visible using light microscopy and reliably revealed the presence of symbionts in host cells. Previous studies [28, 29] showed that in juvenile and adult *Bathymodiolus* mussels, the morphology of epithelial cells colonized by symbionts is remodelled: i) The microvilli that cover all epithelial cells are lost (known as microvillar effacement), and ii) epithelial cells become hypertrophic (swollen) compared to aposymbiotic cells. We identified a third characteristic change in epithelial cells colonized by symbionts that has not received much attention, namely the loss of cilia (Figure 2 c, f and i). We tested if these three morphological characteristics had predictive power for symbiont colonization by analysing 1813 epithelial cells from a subset of six *B. puteoserpentis* individuals (2 pediveliger, 1 plantigrade, 3 post-larvae) and comparing LM images of these cells with their correlated TEM images (average cell count per sample 302). Our analyses showed that all cells predicted to have symbionts in the LM dataset were indeed colonized in the TEM dataset, and likewise, all cells predicted to be aposymbiotic were free of symbionts (Figure S10). We next used our verified morphological characters to reveal the onset of colonization in *B. puteoserpentis* in the subset of six mussel individuals. All pediveliger cells were free of symbionts (*n* = 797 host cells in 2 pediveliger). In plantigrades 15% and in post-larvae 23% of all analysed gill, mantle, foot and retractor muscle epithelia cells were colonized by symbionts (*n* = 336 host cells in 1 plantigrade; *n* = 680 host cells in 3 post-larvae). Finally, we expanded our analysis to LM analyses of serially sectioned individuals, and these confirmed our results from the correlative dataset (all pediveligers were aposymbiotic, n=5; 50% of the plantigrades were colonized by symbionts, n=6; all post-larvae were colonized by symbionts, n=5).

**Figure 2.**
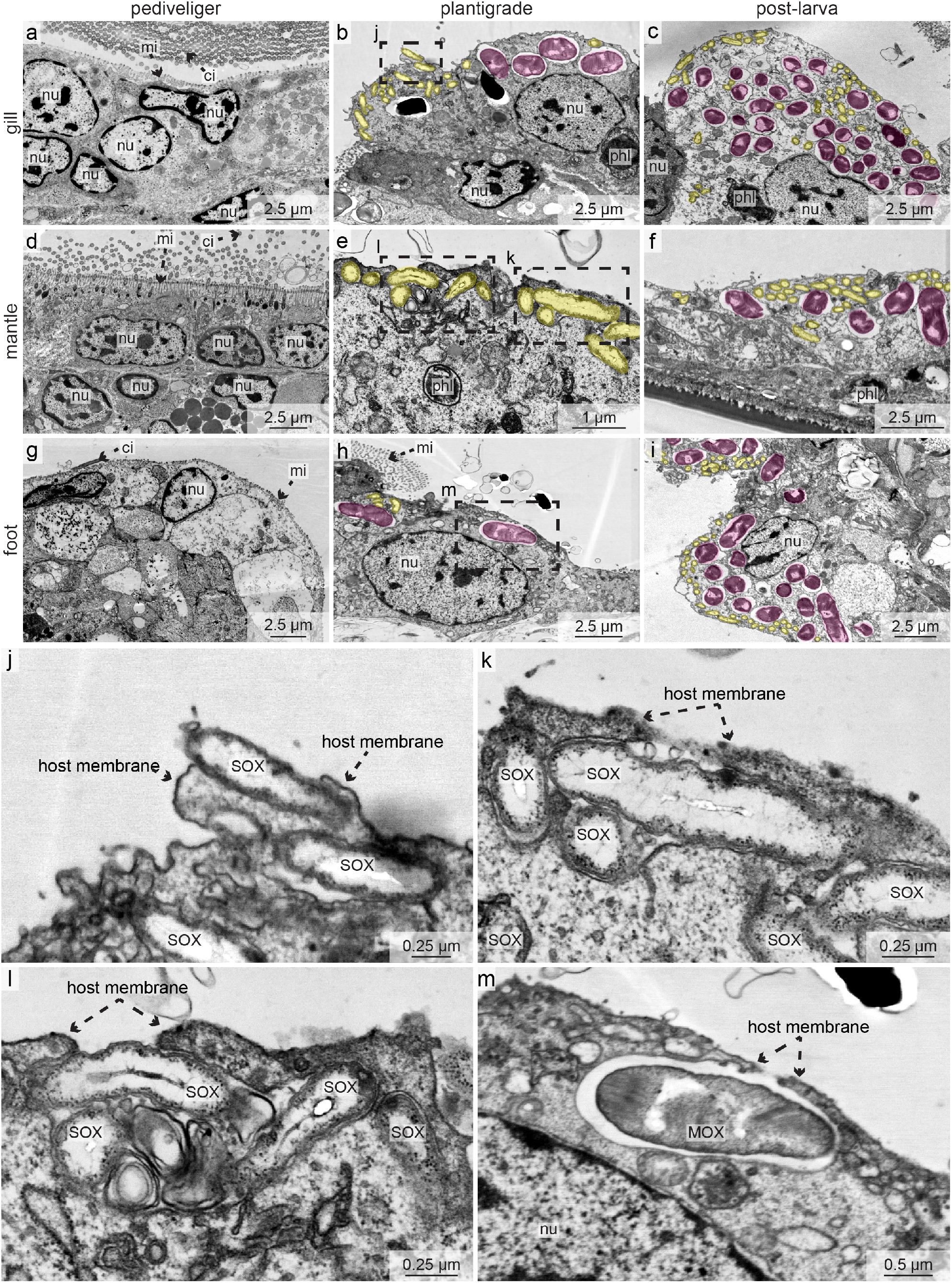
Symbionts first colonize *Bathymodiolus puteoserpentis* at the plantigrade stage. TEM micrographs of gills (**a–c**), mantle (**d–f**) and foot (**g–i**) epithelial tissues of pediveligers (**a, d** and **g**), plantigrades (**b, e** and **h**) and post-larvae (**c, f** and **i**). SOX (yellow) and MOX symbionts (magenta) are highlighted with colour overlays (**a–i**). All epithelial tissues of pediveligers were aposymbiotic (**a, d** and **g**). Colonization by the SOX and MOX symbionts was first observed in the plantigrade stage (**b, e** and **h**). In post-larvae, all epithelial tissues were colonized by both symbiont types (**c, f** and **i**). Dashed boxes indicate regions in which symbionts were in the process of colonizing epithelial tissue (shown magnified in **j – m**). ci, cilia; mi, microvilli; MOX, methane-oxidizing symbiont; nu, nucleus; phl, phagolysosome; SOX, sulphur-oxidizing symbiont. For raw image data see Supplement Figure S14.

Expanding our analyses to all three *Bathymodiolus* species, we found that the pediveligers of all three host species were aposymbiotic (Figure 2 a, d and g). Interestingly, two of the aposymbiotic *B. puteoserpentis* pediveligers had bacterial morphotypes similar to the SOX and MOX symbionts attached to the outside of their shell (Figure S11). The next developmental stage, the plantigrades, were the earliest developmental stage in which we found symbionts in all three host species, with the smallest individuals the plantigrades of *B. puteoserpentis* with a shell length of 432 µm, *B. azoricus* with a shell length of 510 µm and *“B”. childressi* with a shell length of 383 µm. In these plantigrades, we observed symbionts in epithelial cells of the gill filaments, mantle, foot and retractor muscle (Figure 2 b, e, h and j–m). Following symbiont colonization, the gill tissue showed the colonization pattern known from adult animals: the majority of gill cells were bacteriocytes without microvilli or cilia, whereas the only gill cells without symbionts were those at the ventral ends of the gill filaments and at the frontal-to-lateral zones along the length of the filaments, as well as the intercalary cells (Figure S7, S8 and S10). In addition, the epithelial tissues of the mantle, foot and retractor muscles also had symbionts in all three host species (Figure 2 c, f and i). We never observed other bacteria besides the two symbionts in any of the developmental stages, including the intranuclear parasite that infects these mussels, based on FISH analyses with symbiont-specific and eubacterial probes, and TEM analyses of symbiont morphology (Figure 2, Figure 3, Figure S12 and Figure S13).

**Figure 3.**
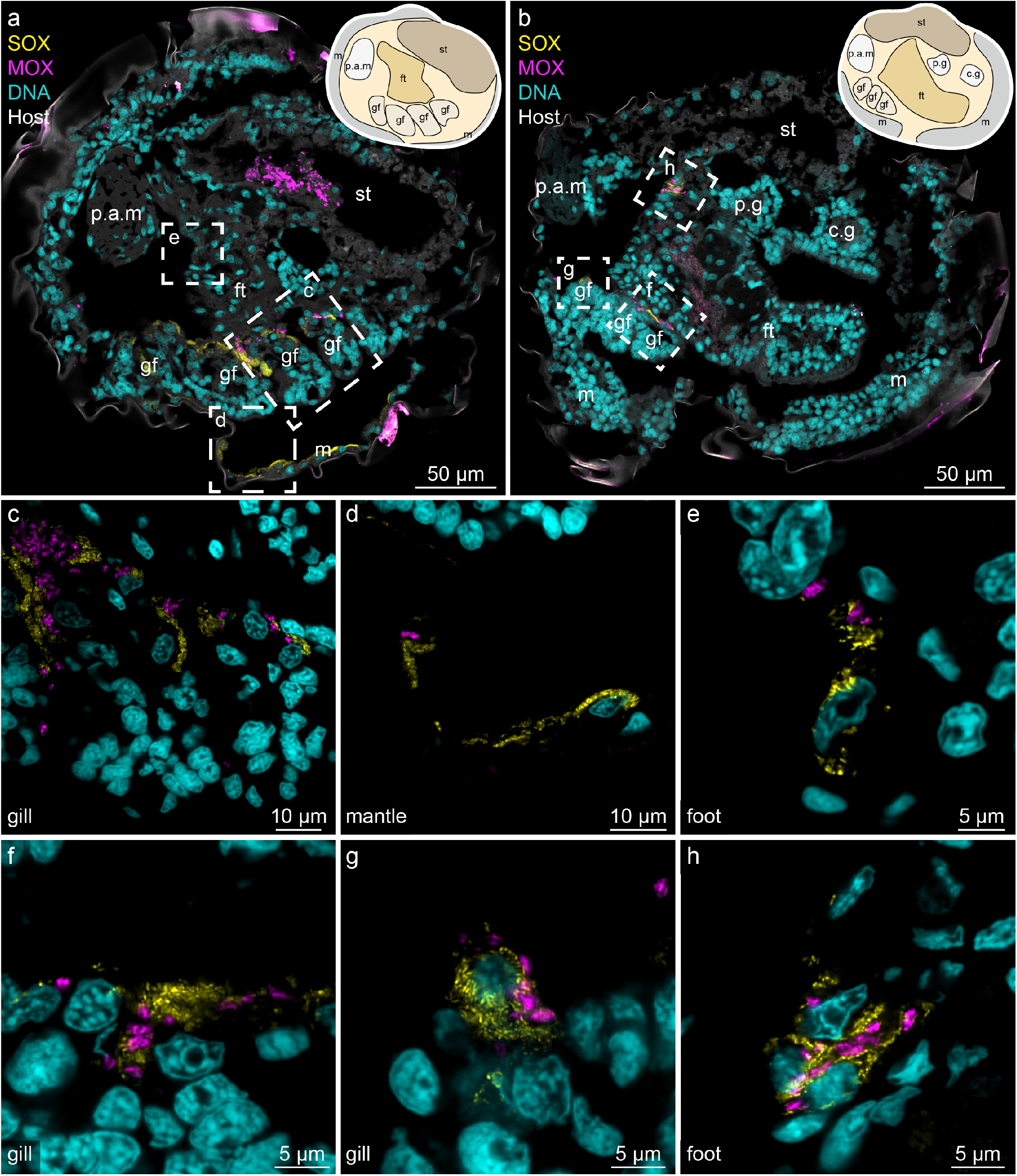
SOX and MOX symbionts colonize all epithelial tissues in *Bathymodiolus puteoserpentis* post-larvae. False-coloured FISH images show probes specific for SOX (yellow, BMARt-193) and MOX symbionts (magenta, BMARm-845) and host nuclei stained with DAPI (cyan). Sagittal cross sections of two different individuals (**a** and **b**) show SOX and MOX symbionts in the gill, foot and mantle epithelia. A schematic drawing of the anatomy is provided in the top right corner for both images. To visualize host tissues, its autofluorescence is shown in grey in **a** and **b**. Dashed boxes indicate magnified regions of the colonized gill, mantle and foot region shown in **c–h**. c.g, cerebral ganglion; ft, foot; m, mantle; gf, gill filament; p.a.m, posterior adductor muscle; p.g, pedal ganglion; st, stomach.

The good preservation of *B. puteoserpentis* specimens allowed us to analyse the process of symbiont colonization in this species. The first visible evidence of symbiont colonization was the presence of only a few bacterial cells in epithelial cells of plantigrades (Figure 2b, e, f, j-m). Bacterial density per gill cell was the lowest in plantigrades with 13.7% of the host cell area occupied by symbionts, but steadily increased in the later developmental stages, reaching up to 29.0% in the post-larval stage and 32.1% in adult mussels (Table S7). With the onset of symbiont colonization, we observed phagolysosomal digestion of the symbionts, in all epithelial cells, even those with only very few symbionts (Figure 2 b – c and e – f). The process of symbiont colonization appeared to be extremely rapid. Nearly all plantigrades had either no symbionts at all, or all of their epithelial tissues were colonized. Only in two out of seven individuals, we occasionally observed SOX and MOX symbionts that were not completely engulfed by the host’s apical cell membrane, which we interpreted as on-going colonization (Figure 2 j – m). These first steps in colonization were particularly common in mantle epithelial cells, while in the same specimens the gill epithelial cells were already fully colonized (Figure 2b, e and f). Furthermore, in the cells where colonization was ongoing, we observed cells that were colonized only by SOX (Figure 2 e), or by both SOX and MOX (Figure 2 h), but we never observed cells that were only colonized by MOX symbionts.

## Discussion

### Post-metamorphosis development in *Bathymodiolus* deviates from the mytilid blueprint

Shell measurements are commonly used to determine the developmental stage of bivalves [14], but our study shows that these are not a reliable characteristic. Our analyses of shell lengths in *Bathymodiolus* developmental stages revealed an overlap in size of 50 µm between pediveligers and plantigrades, as well as between plantigrades and post-larvae. The inconsistency in shell lengths between different developmental stages of *Bathymodiolus* indicates that initiation of metamorphosis is not size dependent and that these mussels are able to delay their metamorphosis while continuing to grow. Such a delay could be due to a lack of settlement cues and/or limited nutrition, similar to what is known from *M. edulis* [30]. Delaying metamorphosis would favour dispersal, potentially leading to an increase in geographic distribution and the colonization of new and remote habitats.

The pre-metamorphosis development of *Bathymodiolus* mussels is very similar to that of their well-studied shallow water relative *M. edulis* [16, 27, 30]. The pediveligers of both *Bathymodiolus* and *Mytilus* have a large velum, foot, mantle epithelium, digestive system, central nerve system and two preliminary gill baskets consisting of three to five gill buds [16]. During the metamorphosis of *Bathymodiolus* mussels, the velum is degraded, organs within the mantle cavity are rearranged and the lipid vesicles in the digestive tract are reduced. In early developmental stages of mussels of the families *Mytilidae* and *Teredinidae*, lipid vesicles serve as storage compounds to sustain the energy-intensive metamorphosis process [16, 31], and they are likely to have a similar function in *Bathymodiolus*. Furthermore, these lipid vesicles could provide energy for supporting movement of the pediveligers during their search for sites to settle [32, 33].

Although early development appears to be conserved between *Bathymodiolus* mussels and their shallow water relatives, marked differences occur as soon as symbiont colonization begins. All colonized epithelial cells lost not only their microvilli but also their cilia, and developed a hypertrophic habitus, as previously documented for bacteriocytes of juveniles and adults [28, 29]. Similar remodelling of the cell surface and induction of the loss of cilia and microvilli has been reported for many bacteria invading a wide range of epithelia [34-36]. Our correlative analyses demonstrate that these cell surface modifications serve as reliable markers for the state of symbiont colonization.

Although it has been known for several decades that the symbionts of *Bathymodiolus* supply them with nutrition, nothing was known about how this affects the development of the digestive system in these hosts. We observed a streamlining of the digestive system after completion of metamorphosis in all three species (Figure 4). The stomach and the intestine straightened and the digestive system changed from the complex looped type found in *Mytilus* to the straight type seen in *Bathymodiolus* adults (Table S8, [37, 38]). Such change in morphology is striking, as in mytilids no further morphological changes occur after metamorphosis Conspicuously, in *Bathymodiolus* this remodelling of the digestive system did not coincide with the first stages of symbiont colonization or metamorphosis, but rather occurred during the transition from post-larvae to the juvenile stage, well after metamorphosis and only when these hosts had become fully colonized by their symbionts (Figure 4). We hypothesize that the streamlining of the digestive system is induced by the shift in nutrition of these hosts. In general, the morphology of an animal’s gastrointestinal tract reflects its food sources. Animals that digest complex foods possess enlarged compartments and lengthened gastrointestinal structures to slow down the flow of digested material and increase the breakdown of complex molecules [39]. We postulate that the first life stages of *Bathymodiolus* rely on filter feeding and require a complex intestine to digest their diverse diet. Once they shift from filter feeding to obtaining most of their nutrition from intracellular symbionts, the morphology of the digestive tract changes. The streamlining of the digestive tract could be an evolutionary strategy to reduce the energy needed for maintaining the digestive tract once the majority of nutrition is covered by intracellular digestion of symbionts in the bacteriocytes. It is interesting to note that adult bathymodiolin mussels of the closely-related genus *Vulcanidas* that occur in shallow waters close to the photic zone (140 m compared to 1000–2500 m depth for the samples studied here) have a pronounced looped intestine [40]. With more food input from the photic zone, filter feeding might play a greater role in the nutrition of *Vulcanidas* mussels, in contrast to the three *Bathymodiolus* species analysed in this study. This raises the question whether streamlining of the digestive tract is a conserved developmental trait among *Bathymodiolus* species, or if the environment and access to external food sources dictates this post-metamorphosis development.

**Figure 4.**
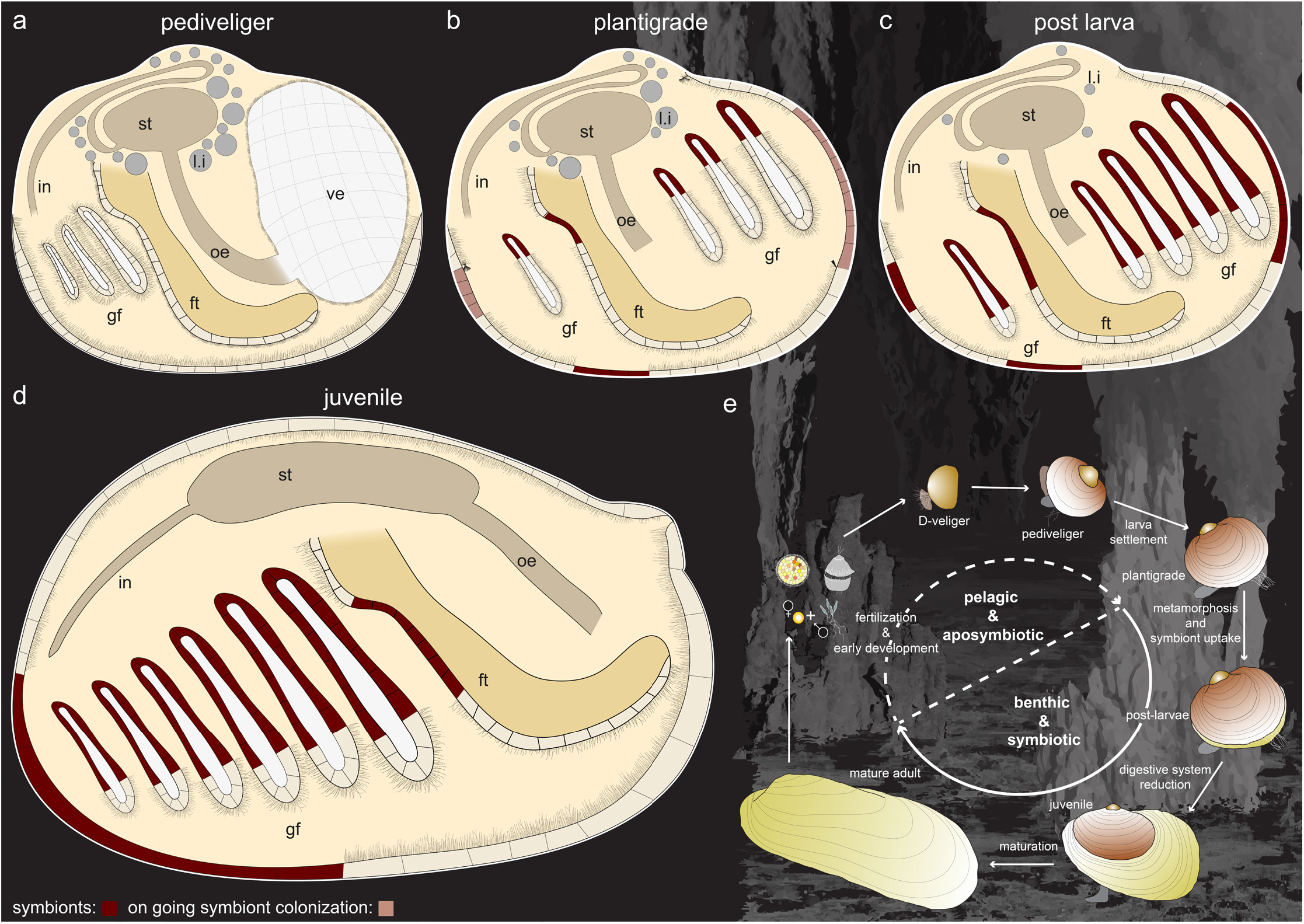
Summary of symbiont colonization and development of bathymodiolin mussels. Pediveliger are aposymbiotic (**a**), and symbiont colonization begins in plantigrades (**b**). By the post-larval (**c**) and juvenile (**d**) stage, all epithelial tissues are fully colonized by symbionts. The digestive system is reduced between the post-larva and juvenile stages, changing from a looped to a straight morphology. (**e**) Schematic of the hypothetical life cycle of bathymodiolin mussels indicating the aposymbiotic pelagic developmental stages and the symbiotic benthic developmental stages. ft, foot; gf, gill filament; in, intestine; l.i, lipid inclusion; oe, oesophagus.

### Symbiont colonization begins during the plantigrade stage as soon as the velum is degraded

As highlighted by Laming, Gaudron [7], how and when *Bathymodiolus* mussels acquire their symbionts has remained, as yet, unclear. Here we used correlative imaging analyses to reveal that the early developmental stages of three *Bathymodiolus* species were aposymbiotic, and narrowed the window of symbiont acquisition down to the plantigrade stage of these hosts (Figure 4). Our analyses revealed that the symbionts colonize the gills only after the velum is lost. As long as the velum is present and active, particles from the surrounding seawater are either transported directly into the digestive system or expelled from the mussel. In *M. edulis*, once the velum is degraded the gill filaments separate further from each other and take over the task of sorting food particles and generating a water current [16, 27]. In the late plantigrade stage of *Bathymodiolus*, we identified a similar increase of space between the gill filaments. It is likely that this morphological change, together with the onset of gill water filtering and particle sorting, is essential for allowing the mussel symbionts to adhere to the gill epithelium and initiate colonization.

The initial colonization of the host by their symbionts appears to be rapid, based on our observation that 22 out of 24 *B. puteoserpentis* individuals were either completely aposymbiotic or fully colonized, and the first stages of symbiont colonization were only visible in two plantigrades. As we were working with preserved samples that represent a snapshot of development, the chance of observing a process depends on how often it occurs and how long it takes. The less frequent or the faster a process happens, the smaller the chance of observing it. We therefore conclude that symbiont colonization occurs rapidly in *Bathymodiolus*, given the small percentage of individuals in which we observed the first steps of colonization. This could reflect the importance for these mussels to quickly acquire symbionts once they have settled because of the lack of energy-rich planktonic nutrition in their environment [41], and the consumption of their internal energy reserves.

In post-larvae and juveniles, the symbionts occur in epithelial cells of the gills, mantle, foot and retractor muscle [14, 42, 43]. Previous work suggested that the symbionts first colonize the mantle epithelia, and from there colonize gill cells as the first gill filaments are formed from mantle tissues [42]. Our data contradicts this assumption, as the gills had already begun to develop in pediveligers before the onset of symbiont colonization. Furthermore, in two *B. puteoserpentis* specimens we detected fully colonized gills, while the mantle tissue was still in the process of being colonized. Our findings indicate that symbionts first colonize gill epithelial cells before colonizing other epithelial tissues, and that the SOX symbionts colonize individual host cells first, before the MOX.

Although all *Bathymodiolus* individuals in this study were collected from mussel beds on the sea floor, the pediveligers were always aposymbiotic. We therefore assume that these early planktonic larval stages were in the process of settling on the sea floor. Recent work has highlighted how bacteria can induce settlement and metamorphosis in marine invertebrates [44]. It is tempting to speculate that the symbiont-like bacteria we observed on the larval shells, or microorganisms in the mussel beds, including possible free-living forms of the symbionts, play a role in inducing the settlement and metamorphosis of *Bathymodiolus* mussels.

Our analyses revealed that *Bathymodiolus* larvae are aposymbiotic during their planktonic phase and first acquire their symbionts when they transition to a benthic lifestyle. This acquisition of symbionts after settlement allows the mussels to recruit locally adapted symbionts. As the geochemistry of vent and seep environments varies strongly across temporal and spatial scales [45], recruiting locally adapted symbiont populations would confer a strong selective advantage to these hosts. Indeed, recent studies have revealed that *Bathymodiolus* mussels host multiple strains of symbionts that vary in key functions, such as the use of energy and nutrient sources, electron acceptors and viral defence mechanisms [46, 47]. By acquiring their symbionts from the sites where they settle, *Bathymodiolus* mussels can establish symbioses with those strains that are best adapted to the local environment.

## Conclusion and outlook

The correlative imaging workflow we used in this study revealed the intricate developmental processes from the subcellular to the whole animal scale that are triggered when deep-sea mussels acquire their symbionts. Our analyses revealed the narrow window in which symbiont acquisition begins and revealed the morphological changes of the digestive system following symbiont uptake. Given that we never observed bacterial morpho- or phylotypes other than the known SOX and MOX symbionts, even in the earliest larval life stages, strong recognition mechanisms must ensure this high specificity. Now that we have identified when and how symbiont colonization takes place in *Bathymodiolus*, a spatial and temporal transcriptomic approach could shed light on the underlying molecular mechanisms of symbiont recognition, acquisition and maintenance, and further our understanding of the interwoven dialogue between animal hosts and their microbial symbionts.

## Data accessibility

LM-data and µCT-data is available on figshare. The DOIs are listed in Table S1.

All supplementary videos are available with the following DOIs: Video S1: https://doi.org/10.6084/m9.figshare.13049765.v2 Video S2: https://doi.org/10.6084/m9.figshare.13049807.v1 Video S3: https://doi.org/10.6084/m9.figshare.13049870.v1 Video S4: https://doi.org/10.6084/m9.figshare.13049882.v1 Video S5: https://doi.org/10.6084/m9.figshare.13049993.v1

## Authors’ contributions

**M.F**., **N.L**. and **N.D**. conceived this study. **M.F**. and **N.L**. wrote the manuscript, with support from **N.D**., and contributions and revisions from all other co-authors. **M.F**. performed the light and fluorescence microscopy, analysed the light, TEM, fluorescence and µCT datasets, reconstructed the 3D models, designed the figures, and produced the videos. **B.G**. helped with the 3D reconstructions and µCT-measurements. **J.U.H**. performed the µCT measurements with help from **B.G**. and **M.F**. and reconstructed the µCT-datasets. **N.L**. did the TEM re-sectioning and measurements.

## Acknowledgements

We thank the captains, crew members and ROV pilots of the cruises M126, M82-3 and NA58. We thank Christian Borowski and Stéphane Hourdez for their valuable contributions to collecting mussel larvae. Furthermore, we gratefully acknowledge Wiebke Ruschmeier for her help in the lab. We would like to thank all those involved in supporting us at the DESY at the P05 beamline of PETRA III (Helmholtz-Zentrum Hereon, Geesthacht, Germany). We would like to thank Benjamin Cooper (Max-Planck Institute for Experimental Medicine, Göttingen) for preliminary sample preparation and Bernhard Ruthensteiner (Zoologische Staatsammlung München) und Frank Melzner (GEOMAR, Kiel) for fruitful discussions.

