## Supplementary Notes, Tables, Figures and Methods for "Coming together – symbiont acquisition and early development of *Bathymodiolus* mussels"

### Content

**Supplementary Note 1 - Detailed morphological description of the *Bathymodiolus* pediveliger**

**Supplementary Note 2 Morphological changes that occur during the metamorphosis**

**Supplementary Methods**

**Supplementary Tables 1-10**

**Supplementary Figures 1-14**

**Supplementary References**

### Supplementary notes

### Supplementary Note 1 - Detailed morphological description of the *Bathymodiolus* pediveliger

#### Velum and mantle

The velum occupied the anterior half of the mantle cavity and the most anterior point was connected to the mantle via a thin membrane near the anterior adductor muscle (Figure S5). The posterior-most part of this membrane was fused with the visceral mass close to the base of the foot (Figure S5). Furthermore, three velar retractor muscles were present. The mantle epithelium consisted of one to two layers of cells (figure 2 d–f, Figure S5, Figure S8 and Figure S7), except for the mantle folds which consisted of multiple cell layers. The folds were located at the most ventral part of the mussels.

#### The foot

The foot occupied 11% of the soft body volume and was anchored to the shell by two pairs of retractor muscles. The posterior pair were anchored to the shell above the posterior adductor muscle and dorsal-ventral at the base of the foot (Figure S5). The anterior retractor muscle pair were located at the base of the foot close to the pedal ganglion and were connected to the shell near the hinge. The surface of the pediveliger foot was either ciliated or covered with microvilli (Figure 2 g). Three glands were present in the foot of *B. puteoserpentis* pediveligers: the white, purple, and byssus glands (Figure S6). The white gland was located at the base of the foot between the pedal ganglion and posterior adductor muscle, and the purple gland was located close to the white gland (Figure S6 a-f). The byssus gland lay on the ventral side of the white gland, and a pair of ciliated ducts merged into the pedal groove at the heel of the foot (Figure S6 b-d).

#### The central nerve system

Pediveliger had three pairs of fully developed ganglia: the cerebral ganglia located dorsally to the oesophagus and close to the apical plate of the velum, the pedal ganglia situated at the base of the foot, and the visceral ganglia located ventrally to the posterior adductor muscle (Figure S5). Each ganglion was fused with its corresponding partner via commissure tissue (Figure S5). The length of the commissure tissues varied between each ganglion pair: longest between the visceral ganglia and shortest between the pedal ganglia. The cerebral, pedal and visceral ganglia were connected via connective tissues, all of which were fully developed (Figure S5). The cerebro-visceral and cerebro-pedal connectives left the posterior region of each cerebral ganglion as one entity, which then split near the region of the larval foot. On each side, the cerebro-pedal connective dove ventrally towards its respective pedal ganglion, while each cerebro-visceral connective extended posteriorly towards its respective visceral ganglion.

#### The digestive system

The *Bathymodiolus* pediveliger possessed a fully developed digestive system, composed of mouth, oral labial palp, oesophagus, stomach, two digestive glands, the style sac with crystalline style, the gastric shield, an s-shaped looped intestine and a mid-gut. It was located in the most dorsal part of the mussel beneath the hinge and occupied 29.4% of the soft tissue volume. The oral labial palp was located at the posterior end of the velum near the base of the foot. It was connected via a thin membrane to the velum (Figure S4). No primary topography (rafts) was visible in any of the pediveliger oral labial palps, in contrast to the labial palps of adult mussels. The epithelial cells of the oral labial palps were ciliated and had microvilli. The oesophagus began at the posterior edge of the velum and led into the stomach. Most of the oesophagus was located perpendicular to the hinge line beneath the foot (Figure S5). The cells lining the oesophagus were ciliated and had microvilli. In the pediveliger stage, the stomach was the largest organ of the digestive system. It was situated in the middle of the mussel, underneath the base of the foot and the pedal ganglion (Figure S5). The style sac was located at the posterior end of the stomach and had a diameter of 30–40 µm. The cells lining the style sac were densely ciliated and had large vacuoles in their cytoplasm (Figure S5). The crystalline style was observed in pediveliger stages but not in every specimen (probably due to sample preservation). Membrane-bound lipid vesicles were found in the digestive cells of the stomach and had a mean diameter of 15.85 µm ($n$ = 75 µm). In the earliest pediveliger, these lipid vesicles formed up to 12.8% of the entire soft body volume (Figure S5). The gastric shield was found at the dorsal wall of the stomach (Figure S5) and its cells were densely ciliated. The mid gut left the stomach at the ventral wall anterior to the style sac. It passed backwards and turned dorsally to fuse into the intestine, which then passed towards the anal papilla. The part of the digestive tract containing the mid gut and the intestine looped in an s-shape dorsally under the stomach (Figure S5). The ciliated anal papilla was located close to the posterior adductor muscle (Figure S5).

#### The gill basket

In the *Bathymodiolus* pediveliger stage, each gill basket consisted of rudimentary gill filaments and the developing budding zone. The budding zone is a meristem-like zone, which in later developmental stages continuously generates new gill filaments at the posterior end of the gill basket. In mytilids, the first three gill filaments are developed almost simultaneously [1] out of the mantle epithelia, and all subsequent filaments are developed by the budding zone. The shortest filaments were found at the most posterior end of the descending lamellae next to the budding zone (Figure 1 a and Figure S5). The pediveliger gill basket had the fewest rudimentary gill filaments in comparison with later developmental stages (3-4 in *B. puteoserpentis* and 5 in *B. azoricus* and *"B". childressi*; Figure S2). Furthermore, these gill filaments were the shortest dorso-ventrally from the gill axis to the ciliated ends (mean length 81 µm, SD = 10 µm, $n$ = 16 in *B. puteoserpentis*, mean length 97 µm, SD = 18 µm, $n$ = 10 in *"B". childressi,* and mean length 78 µm, SD = 12 µm, $n$ = 10 in *B. azoricus*). The gill axis (dorsal tissue that supports the gill lamellae) was parallel to the hinge line (figure 1 a). The space between individual gill filaments of one gill basket ranged from 9 µm to 27 µm in the *B. puteoserpentis* pediveliger (Figure S5). Correlative light and electron microscopy showed that the surface of the entire gill filament was densely covered with cilia and microvilli (figure 2 a) and no symbiotic bacteria were present.

### Supplementary Note 2 Morphological changes that occur during the metamorphosis

#### The digestive system

The *Bathymodiolus* pediveliger had a fully developed digestive system with a looped mid gut and intestine (Figure S 9 a-e). The rafts of the labial palps were first observed in a post-larva with a size of 436 µm. During the metamorphosis, the apical plate moved into the oral labial palp, which then split into the upper and lower labial palp (Figure S 5, S 7 and S 8). Instead of being located beneath the foot and parallel to the hinge line, the post-larval oesophagus had a more perpendicular position (Figure S8). Organs like the stomach, oesophagus, intestine and style sac increased in size but retained their general morphological traits (Figure 1, Figure S5, S7 and S8). During the transition from a post larvae to a juvenile mussel, the complex digestive system got streamlined and the looped intestine straightened. Furthermore, the volume of lipid droplets decreased from 12.83% of the soft-body volume in pre-metamorphosis mussels to 1.8% in post-metamorphosis mussels, and to their complete absence in juveniles (Figure 1, Figure S5, S7 and S8, and Table S6).

#### Gill development and growth

During metamorphosis, the gill filament number increased by one in all species (*B. puteoserpentis,* 5; *B. azoricus,* 6; and *"B". childressi,* 6). Furthermore, reorientation of the gill basket in the mantle cavity led to the first gill filament pair (which are attached to the visceral mass along their entire anterior face) being located closer to the mouth and the labial palps. Depending on the specimen, the posterior end of the gill basket was tilted by 20–40° (figure 1a to c). This reorientation aligned the growth axis of the gill with the length axis of the mussel. In the post-larval stage, there was an increase in the dorso-ventral filament length (increasing from 68 µm to 159 µm in *B. puteoserpentis,* from 73 µm to 215 µm in *"B". childressi* and from 68 µm 110 µm in *B. azoricus*) and the frontal-abfrontal filament depth. Gill filaments of *"B". childressi* (mean = 125.49 µm, sd = 33.77 µm, *n* = 52) were larger than *B. puteoserpentis* (mean = 115.21 µm, sd = 24.56 µm, *n* = 40) and *B. azoricus* (mean = 93.58 µm, sd = 18.17 µm, *n* = 10). Additionally, gill filaments separated from each other and the gaps between the filaments increased (up to 120 µm). The gill filament tips disconnected from each other after metamorphosis (figure 1 c and Figure S7). At the end of metamorphosis, a distinct ciliated and non-ciliated region could be identified (figure S8). The non-ciliated region was filled with bacteria (figure 2 c). This symbiont colonization led to the hypertrophic morphology of the epithelial cells. The hypertrophy contributed to a drastic increase of the relative gill volume from 4.6% to 28% after the mussels were colonized (Table S6).

In one 2-mm *B. puteoserpentis* juvenile mussel a well-developed right and left gill could be identified. Each gill consisted of the descending (25 gill filaments) and ascending lamella (9 gill filaments) of the inner demibranch. In one 3-mm mussel the descending lamella (4 gill filaments) of the outer demibranch was in the process of development. Upon reaching 1 cm shell length, the gill was completely developed and consisted of the inner and outer demibranch with the ascending and descending lamellae.

### Supplementary Methods

#### Decalcification

For decalcification, mussels were re-hydrated with an ethanol series of decreasing concentration (60% (v/v) to 0%, in steps of 10%), with each step lasting 10 min. The samples were washed twice with 1x phosphate-buffered saline (PBS) and transferred into a 0.5 M Ethylenediaminetetraacetic acid (EDTA) PBS solution. Decalcification was performed for 24–240 h depending on the size of the specimens. Samples stored in PHEM-buffer were not treated with EDTA because PHEM-buffer includes EGTA, which decalcifies similarly to EDTA.

#### µCT measurements

The two adult *Bathymodiolus* mussels, *B. childressi* (H1425-019-J1) and *B. azoricus* (D4MS) were scanned with a laboratory-based µCT at the Zoologische Staatssammlung in Munich, Germany. Both mussels were stained in phosphotungstic acid, dissolved in ethanol and scanned in solution as described in [2].

#### Embedding, sectioning and staining for histological analysis

All glutaraldehyde-fixed samples were post fixed with 1% (v/v) osmium tetroxide ($OsO_4$) for 1-2 h at 4 °C and washed three times with PHEM. The mussels were dehydrated with an ethanol series of increasing concentration (30% (v/v), 50% (v/v), 70% (v/v), 80% (v/v), 90% (v/v) and 100% (v/v)) at -10 °C, with each step lasting 10 min. The tissue was then transferred into a 50:50 mixture of ethanol and acetone, followed by 100% acetone. Samples were infiltrated with low-viscosity resin (Agar Scientific, UK) step-wise with a 1:3 resin:acetone mixture for 2 h, followed by 2 h in a 1:1 solution, and overnight in a 3:1 solution. Samples were then transferred twice into pure resin for 2 h. The samples were polymerized at 60–65 °C for 48 h.

#### Double-labelled oligonucleotides -FISH

In this study, double-labelled oligonucleotides (Dope-FISH) were used for the *in situ* hybridization. Hybridization was performed on whole mussel sections of *B. puteoserpentis*. Sections were baked in a vertical position for 1 h at 60 °C to improve adhesion to the slides. Slides were de-waxed and rehydrated by immersing them three times in 100% (v/v) RotiHistol for 10 min each, followed by a decreasing ethanol series (96% (v/v), 80% (v/v), 70% (v/v) and 50% (v/v), 10 min each). Sections were air dried and encircled with a liquid blocker (PAP-Pen, Science Services). For *in situ* hybridization, general probes targeting conserved regions of the 16S rRNA in the domain Bacteria, as well as specific probes for targeting SOX and MOX bacteria were used. As a negative control, an oligonucleotide (non-338) labelled with Atto-550 was used, complementary to the probe EUB I-III, to monitor nonspecific binding (Table S3). Depending on the section size, 20–300 µl of hybridization mixture was applied per section. The 8.4 pmol probe stock solution was diluted 1:10 with the hybridization buffer (Table S9), which contained 30–35% (v/v) of formamide, before the mixture was applied onto the sections. The hybridizations were performed at 46 °C for 3 h in hybridization chambers. To prevent changes in concentration of the hybridization solution through evaporation, a tissue was saturated with 2 ml of the 30–35% formamide solution and placed in the chamber, thereby assuring equilibrium between the hybridization solution and the surrounding gas phase. After hybridization, samples were washed for 15 min in pre-warmed (48 °C) corresponding washing buffer (Table S10) and then dipped once in ddH2O. DAPI was applied as a nucleic acid counter-stain. The slides were incubated for 10 min at room temperature, washed twice in ddH2O and dried.

**3D reconstruction**

Prior to the 3D reconstructions, LM-images were stitched and aligned in *x*, *y* and *z* direction with the software Fiji ver. V1.52p and the plugin TrackEM2, and exported. Image files were imported using “import sequences as grid”. Stitching and blending of the layers was performed choosing “montage multiple layers”. For the automated alignment, a linear alignment was used, allowing for translation and rotation. Individual misalignments were manually corrected using the “aligning with landmarks” option. Exported LM image stacks were used for 3D surface reconstructions.

3D surface reconstructions were performed in Amira 6.7.0 (ThermoFisher Scientific). Semi-automated (via thresholds) and manual segmentation were used to label individual organs. The individual labels for each organ were separated with the *arithmetic tool* function: A == 1 (no. of material) and rendered into a 3D surface model using the following steps: *create surface*, *reduce faces*, *remesh surface* and *smooth surface*. *Faces* were reduced down to a value of 50,000–100,000. Surfaces were remeshed and the percentage value in the ‘Desired Size’ port was kept at 50%. All surface meshes were visualized with the Direct Normals shading mode under *Surface View*.

Co-registration of SRμCT, LM and TEM data was carried out following [3]. For visualisation, LM and TEM data were displayed simultaneously as orthographic slices in a single AMIRA 3D scenario. For the selection of virtual planes within the data sets, the slice tool was used. Individual TEM images were co-registered based on their corresponding LM image, using Adobe Photoshop CS5 (Adobe Systems Software Ireland Ltd.).

### Supplementary tables

**Table S1. Overview of samples used for the morphological analyses , SRµCT, µCT, FISH and TEM.** GA, glutaraldehyde; PFA, paraformaldehyde.

| species | sample identifier | developmen-tal stage | shell length (µm) | shell height (µm) | sampling location | latitude | longitude | sampling depth (m) | sam-pling year | analyses | fixation | storage buffer | status of symbi-ont colonization | DOI |
|---|---|---|---|---|---|---|---|---|---|---|---|---|---|---|
| *B. puteoser-pentis* | Bp-hpf | plantigrade | 372.31 | 375.66 | Semenov-2 | 13.513 N | 44.962 W | -2446.5 | 2016 | histology / TEM | 2.5% GA | PHEM | apo- symbiotic | |
| | 1555-30 | post-larva | 372.42 | 353.1 | Semenov-2 | 13.513 N | 44.962 W | -2446.5 | 2016 | SRµCT / histology | 2.5% GA | PHEM | symbiotic | |
| | 1555-27 | plantigrade | 385.47 | 417.85 | Semenov-2 | 13.513 N | 44.962 W | -2446.5 | 2016 | SRµCT / histology | 2.5% GA | PHEM | apo- symbiotic | 10.6084/m9. figshare.14247488.v1 |
| | 1555-16 | pediveliger | 399.37 | 420.37 | Semenov-2 | 13.513 N | 44.962 W | -2446.5 | 2016 | histology | 2.5% GA | PHEM | apo- symbiotic | 10.6084/m9. figshare.14247395 |
| | 1555-06 | pediveliger | 404.99 | 427.36 | Semenov-2 | 13.513 N | 44.962 W | -2446.5 | 2016 | histology / TEM | 2.5% GA | PHEM | apo- symbiotic | 10.6084/m9. figshare.14247329.v1 |
| | 1556-61 | pediveliger | 407.91 | 382.31 | Semenov-2 | 13.513 N | 44.962 W | -2446.5 | 2016 | Dope-FISH | 2% PFA | PBS / ethanol | apo- symbiotic | |
| | 1555-31 | pediveliger | 409.3 | 408.16 | Semenov-2 | 13.513 N | 44.962 W | -2446.5 | 2016 | histology | 2.5% GA | PHEM | apo- symbiotic | 10.6084/m9. figshare.14247980.v1 |
| | 1555-28 | pediveliger | 409.34 | 415.8 | Semenov-2 | 13.513 N | 44.962 W | -2446.5 | 2016 | SRµCT / histology | 2.5% GA | PHEM | apo- symbiotic | 10.6084/m9. figshare.14247698.v1 |
| | 1556-26 | plantigrade | 410.38 | 428.54 | Semenov-2 | 13.513 N | 44.962 W | -2446.5 | 2016 | Dope-FISH | 2% PFA | PBS / ethanol | only SOX | |
| | 1555-45 | plantigrade | 411.13 | 442.17 | Semenov-2 | 13.513 N | 44.962 W | -2446.5 | 2016 | histology | 2.5% GA | PHEM | apo- symbiotic | 10.6084/m9. figshare.14248103.v1 |
| | 1555-04 | pediveliger | 411.23 | 402.91 | Semenov-2 | 13.513 N | 44.962 W | -2446.5 | 2016 | histology / TEM | 2.5% GA | PHEM | apo- symbiotic | 10.6084/m9. figshare.12367259 |
| | 1556-29 | post-larva | 414.18 | 394.26 | Semenov-2 | 13.513 N | 44.962 W | -2446.5 | 2016 | Dope-FISH | 2% PFA | PBS / ethanol | only SOX | |
| | 1555-14 | post-larva | 414.32 | 394.93 | Semenov-2 | 13.513 N | 44.962 W | -2446.5 | 2016 | SRµCT / histology | 2.5% GA | PHEM | symbiotic | |
| | 1555-42 | plantigrade | 417.82 | 413.34 | Semenov-2 | 13.513 N | 44.962 W | -2446.5 | 2016 | histology | 2.5% GA | PHEM | apo- symbiotic | 10.6084/m9. figshare.14248046.v1 |
| | 1556-05 | pediveliger | 418.47 | 463.48 | Semenov-2 | 13.513 N | 44.962 W | -2446.5 | 2016 | Dope-FISH | 2% PFA | PBS / ethanol | apo- symbiotic | 10.6084/m9. figshare.12367247 |
| | 1555-03 | post-larva | 432.29 | 447.89 | Semenov-2 | 13.513 N | 44.962 W | -2446.5 | 2016 | histology / TEM | 2.5% GA | PHEM | symbiotic | 10.6084/m9. figshare.14247305.v3 |
| | 1556-62 | post-larva | 432.8 | 433.97 | Semenov-2 | 13.513 N | 44.962 W | -2446.5 | 2016 | Dope-FISH | 2% PFA | PBS / ethanol | symbiotic | |
| | 1556-07 | post-larva | 434.02 | 445.35 | Semenov-2 | 13.513 N | 44.962 W | -2446.5 | 2016 | Dope-FISH | 2% PFA | PBS / ethanol | symbiotic | |
| | 1555-01 | post-larva | 435.53 | 404.12 | Semenov-2 | 13.513 N | 44.962 W | -2446.5 | 2016 | histology / TEM | 2.5% GA | PHEM | symbiotic | 10.6084/m9. figshare.11709399 |
| | 1555-05 | plantigrade | 436.44 | 390.65 | Semenov-2 | 13.513 N | 44.962 W | -2446.5 | 2016 | histology / TEM | 2.5% GA | PHEM | start of coloni-zation | 10.6084/m9. figshare.12367247.v1 |
| | 1555-02 | post-larva | 439.64 | 493.34 | Semenov-2 | 13.513 N | 44.962 W | -2446.5 | 2016 | histology / TEM | 2.5% GA | PHEM | symbiotic | 10.6084/m9. figshare.14248436.v1 |
| | 1556-49 | juvenile | 2053.09 | 1451.25 | Semenov-2 | 13.513 N | 44.962 W | -2446.5 | 2016 | Dope-FISH | 2% PFA | PBS / ethanol | symbiotic | |
| | 1555-47 | juvenile | 2290.09 | 1508.85 | Semenov-2 | 13.513 N | 44.962 W | -2446.5 | 2016 | SRµCT | 2.5% GA | PHEM | symbiotic | 10.6084/m9. figshare.14248496.v1 |
| | 1555-22 | juvenile | 3072.61 | 2060.11 | Semenov-2 | 13.513 N | 44.962 W | -2446.5 | 2016 | SRµCT / histology | 2.5% GA | PHEM | symbiotic | 10.6084/m9. figshare.12728075.v1 |
| *“B”. childressi* | M102 | plantigrade | 383.82 | 406.78 | Mississippi Canyon 853 | 28.123 N | -89.139 E | -1071.00 | 2015 | histology | 2.5% GA | PHEM | start of coloni-zation | 10.6084/m9. figshare.14248553.v1 |
| | Bc-hpf-small | plantigrade | 389.94 | 406.16 | Mississippi Canyon 853 | 28.123 N | -89.139 E | -1071.00 | 2015 | histology | 2.5% GA | PHEM | start of coloni-zation | 10.6084/m9. figshare.14248670.v1 |

| | | | | | | | | | | | | | | |
|---|---|---|---|---|---|---|---|---|---|---|---|---|---|---|
| | Bc-hpf-large | post-larva | 429.08 | 500.32 | Mississippi Canyon 853 | 28.123 N | -89.139 E | -1071.00 | 2015 | histology | 2.5% GA | PHEM | symbiotic | 10.6084/m9. figshare.14248643.v1 |
| | M103 | post-larva | 433.13 | 463.17 | Mississippi Canyon 853 | 28.123 N | -89.139 E | -1071.00 | 2015 | histology | 2.5% GA | PHEM | symbiotic | 10.6084/m9. figshare.14248559.v1 |
| | M101 | pediveliger | 434.37 | 447.76 | Mississippi Canyon 853 | 28.123 N | -89.139 E | -1071.00 | 2015 | SRµCT / his-tology / TEM | 2.5% GA | PHEM | apo- symbiotic | 10.6084/m9. figshare.14248589.v1 |
| | H1425-019-J1 | adult | 56310 | 31740 | Green Can-yon 234 | 27.746 N | -91.222 E | -540.00 | 2015 | µCT | David-son's fixative | PBS / ethanol | symbiotic | 10.6084/m9. figshare.5458240.v1 |
| *B. azoricus* | 1643-03 | pediveliger | 452.23 | 483.72 | Bubbylon | 37.801 N | -31.537 E | -1002.00 | 2010 | histology / TEM | 2.5% GA | Trump's solution | apo- symbiotic | |
| | 1624-13 | pediveliger | 465.12 | 522.93 | Bubbylon | 37.801 N | -31.537 E | -1002.00 | 2010 | histology | 2% PFA | PBS / ethanol | apo- symbiotic | 10.6084/m9. figshare.14248910.v1 |
| | 1643-02 | post-larva | 498.16 | 510.22 | Bubbylon | 37.801 N | -31.537 E | -1002.00 | 2010 | SRµCT / histology | 2.5% GA | Trump's solution | symbiotic | |
| | 1624-17 | post-larva | 549.1 | 583.44 | Bubbylon | 37.801 N | -31.537 E | -1002.00 | 2010 | histology | 2% PFA | PBS / ethanol | symbiotic | 10.6084/m9. figshare.14249006 |
| | 1643-01 | plantigrade | 566.76 | 597.38 | Bubbylon | 37.801 N | -31.537 E | -1002.00 | 2010 | histology | 2.5% GA | Trump's solution | start of coloni-zation | 10.6084/m9. figshare.14249021.v1 |
| | 1626-01 | juvenile | 2170 | 1430 | Bubbylon | 37.801 N | -31.537 E | -1002.00 | 2010 | SRµCT | 2% PFA | PBS / ethanol | symbiotic | |
| | 1629-124 | juvenile | 2600 | 1500 | Bubbylon | 37.801 N | -31.537 E | -1002.00 | 2010 | SRµCT | 2% PFA | PBS / ethanol | symbiotic | |
| | 1624-121 | adult | 10600 | 6000 | Bubbylon | 37.801 N | -31.537 E | -1002.00 | 2010 | SRµCT | 2% PFA | PBS / ethanol | symbiotic | |
| | D4MS | adult | 53352 | 17078 | Montsegur | 37.288 N | -32.276 E | -1700.00 | 2013 | µCT | 2% PFA | PBS / ethanol | symbiotic | 10.6084/m9. figshare.5458234.v1 |
| *M. edulis* | 252-1 | juvenile | 2198 | 1480 | Kiel | 54.394 N | 10.190 E | -1.5 | 2018 | SRµCT | 2.5% GA | PHEM | aposymbiotic | |

**Table S2. Shell dimensions of *Bathymodiolus* individuals.** Individuals are sorted by increasing shell length. *B. azoricus* mussels were sampled at the Bubbylon vent at 1002 m water depth, *B. puteoserpentis* at the Semenov-2 vent at 2446 m water depth and *"B". childressi* mussels at the Mississiooi Canyon 853 at 1071 m water depth. Individuals are sorted by increasing shell length.

| species | identifier | developmental stage | shell length (µm) | shell height (µm) |
|---|---|---|---|---|
| *B. azoricus* | 1626-12 | pediveliger / plantigrade | 392.36 | 505.18 |
| *B. azoricus* | 1624-45 | pediveliger / plantigrade | 438.55 | 526.14 |
| *B. azoricus* | 1624-41 | pediveliger / plantigrade | 448.79 | 523.95 |
| *B. azoricus* | 1643-03 | pediveliger | 452.23 | 483.72 |
| *B. azoricus* | 1626-15 | pediveliger / plantigrade | 459.18 | 501.96 |
| *B. azoricus* | 1624-12 | pediveliger / plantigrade | 460.77 | 516.14 |
| *B. azoricus* | 1626-05 | pediveliger / plantigrade | 464.71 | 466.27 |
| *B. azoricus* | 1624-13 | pediveliger | 465.12 | 522.93 |
| *B. azoricus* | 1626-32 | pediveliger / plantigrade | 466.08 | 530.28 |
| *B. azoricus* | 1624-02 | pediveliger / plantigrade | 470.00 | 510.00 |
| *B. azoricus* | 1624-29 | pediveliger / plantigrade | 470.86 | 525.05 |
| *B. azoricus* | 1624-56 | post-larva | 472.50 | 540.31 |
| *B. azoricus* | 1626-21 | pediveliger / plantigrade | 474.14 | 554.33 |
| *B. azoricus* | 1626-33 | post-larva | 477.36 | 514.94 |
| *B. azoricus* | 1626-19 | pediveliger / plantigrade | 477.45 | 509.89 |
| *B. azoricus* | 1624-36 | post-larva | 480.93 | 514.35 |
| *B. azoricus* | 1624-25 | post-larva | 480.95 | 524.56 |
| *B. azoricus* | 1624-38 | pediveliger / plantigrade | 481.45 | 516.30 |
| *B. azoricus* | 1624-06 | pediveliger / plantigrade | 482.46 | 529.78 |
| *B. azoricus* | 1626-30 | pediveliger / plantigrade | 485.83 | 542.96 |
| *B. azoricus* | 1624-24 | pediveliger / plantigrade | 486.43 | 543.35 |
| *B. azoricus* | 1624-28 | post-larva | 486.57 | 522.45 |
| *B. azoricus* | 1626-10 | post-larva | 487.00 | 534.74 |
| *B. azoricus* | 1626-07 | pediveliger / plantigrade | 488.52 | 515.81 |
| *B. azoricus* | 1626-11 | pediveliger / plantigrade | 490.03 | 545.64 |
| *B. azoricus* | 1624-31 | post-larva | 490.71 | 548.67 |
| *B. azoricus* | 1624-03 | post-larva | 491.18 | 549.80 |
| *B. azoricus* | 1626-06 | pediveliger / plantigrade | 496.85 | 538.00 |
| *B. azoricus* | 1626-14 | post-larva | 497.18 | 553.92 |
| *B. azoricus* | 1643-02 | post-larva | 498.16 | 510.22 |
| *B. azoricus* | 1624-04 | pediveliger / plantigrade | 502.44 | 513.91 |
| *B. azoricus* | 1626-08 | pediveliger / plantigrade | 502.87 | 515.49 |
| *B. azoricus* | 1624-33 | post-larva | 503.23 | 545.05 |
| *B. azoricus* | 1624-19 | post-larva | 505.57 | 524.99 |
| *B. azoricus* | 1624-32 | pediveliger / plantigrade | 506.68 | 520.30 |
| *B. azoricus* | 1626-29 | pediveliger / plantigrade | 508.28 | 527.29 |
| *B. azoricus* | 1624-52 | post-larva | 508.98 | 522.91 |
| *B. azoricus* | 1624-50 | post-larva | 509.09 | 526.21 |
| *B. azoricus* | 1624-51 | post-larva | 509.93 | 555.65 |
| *B. azoricus* | 1624-01 | post-larva | 510.00 | 540.00 |
| *B. azoricus* | 1624-05 | pediveliger / plantigrade | 511.57 | 538.75 |
| *B. azoricus* | 1626-13 | post-larva | 512.08 | 557.19 |
| *B. azoricus* | 1626-34 | pediveliger / plantigrade | 512.93 | 496.23 |
| *B. azoricus* | 1624-55 | post-larva | 517.08 | 546.18 |
| *B. azoricus* | 1626-23 | pediveliger / plantigrade | 518.94 | 524.56 |
| *B. azoricus* | 1626-24 | pediveliger / plantigrade | 523.11 | 547.44 |
| *B. azoricus* | 1624-20 | post-larva | 523.27 | 547.95 |
| *B. azoricus* | 1626-16 | pediveliger / plantigrade | 525.34 | 566.51 |
| *B. azoricus* | 1624-61 | post-larva | 525.89 | 549.88 |
| *B. azoricus* | 1624-09 | pediveliger / plantigrade | 529.03 | 556.03 |
| *B. azoricus* | 1624-43 | post-larva | 535.11 | 552.63 |

| | | | | |
|---|---|---|---|---|
| *B. azoricus* | 1624-22 | post-larva | 538.01 | 522.93 |
| *B. azoricus* | 1624-10 | pediveliger / plantigrade | 539.56 | 571.65 |
| *B. azoricus* | 1624-62 | post-larva | 548.75 | 578.23 |
| *B. azoricus* | 1624-17 | post-larva | 549.10 | 583.44 |
| *B. azoricus* | 1624-08 | post-larva | 549.94 | 569.43 |
| *B. azoricus* | 1624-23 | post-larva | 564.93 | 552.87 |
| *B. azoricus* | 1624-44 | post-larva | 566.20 | 507.37 |
| *B. azoricus* | 1643-01 | plantigrade | 566.76 | 597.38 |
| *B. azoricus* | 1624-48 | post-larva | 575.36 | 562.86 |
| *B. azoricus* | 1624-47 | post-larva | 578.06 | 581.35 |
| *B. azoricus* | 1624-63 | post-larva | 583.08 | 605.97 |
| *B. azoricus* | 1624-30 | post-larva | 584.02 | 568.64 |
| *B. azoricus* | 1624-54 | post-larva | 592.51 | 530.33 |
| *B. azoricus* | 1624-40 | post-larva | 596.56 | 615.77 |
| *B. azoricus* | 1624-34 | post-larva | 599.79 | 614.15 |
| *B. azoricus* | 1626-28 | post-larva | 602.94 | 559.04 |
| *B. azoricus* | 1624-58 | post-larva | 611.63 | 579.10 |
| *B. azoricus* | 1624-46 | post-larva | 613.10 | 623.03 |
| *B. azoricus* | 1626-26 | post-larva | 614.86 | 542.38 |
| *B. azoricus* | 1624-07 | post-larva | 616.61 | 592.04 |
| *B. azoricus* | 1626-25 | post-larva | 624.21 | 562.41 |
| *B. azoricus* | 1624-26 | post-larva | 631.25 | 550.14 |
| *B. azoricus* | 1626-31 | post-larva | 638.35 | 561.34 |
| *B. azoricus* | 1626-09 | post-larva | 640.41 | 567.16 |
| *B. azoricus* | 1624-49 | post-larva | 643.33 | 606.26 |
| *B. azoricus* | 1624-64 | post-larva | 643.73 | 618.60 |
| *B. azoricus* | 1624-11 | post-larva | 658.55 | 632.35 |
| *B. azoricus* | 1624-35 | post-larva | 665.63 | 620.81 |
| *B. azoricus* | 1624-59 | post-larva | 672.76 | 630.40 |
| *B. azoricus* | 1624-14 | post-larva | 690.20 | 619.74 |
| *B. azoricus* | 1624-18 | post-larva | 706.98 | 681.92 |
| *B. azoricus* | 1624-65 | post-larva | 707.07 | 681.16 |
| *B. azoricus* | 1626-17 | post-larva | 717.93 | 683.47 |
| *B. azoricus* | 1629-122 | post-larva | 730.00 | 661.00 |
| *B. azoricus* | 1624-21 | post-larva | 731.39 | 646.22 |
| *B. azoricus* | 1626-18 | post-larva | 740.99 | 651.74 |
| *B. azoricus* | 1624-60 | post-larva | 744.97 | 588.15 |
| *B. azoricus* | 1626-22 | post-larva | 754.09 | 679.86 |
| *B. azoricus* | 1629-03 | post-larva | 757.01 | 809.73 |
| *B. azoricus* | 1626-20 | post-larva | 772.60 | 644.87 |
| *B. azoricus* | 1626-27 | post-larva | 794.91 | 662.61 |
| *B. azoricus* | 1624-57 | post-larva | 816.73 | 676.32 |
| *B. azoricus* | 1629-02 | post-larva | 833.91 | 854.09 |
| *B. azoricus* | 1624-39 | post-larva | 842.26 | 739.78 |
| *B. azoricus* | 1629-01 | post-larva | 948.74 | 1000.00 |
| *B. azoricus* | 1624-27 | post-larva | 1005.83 | 788.64 |
| *B. azoricus* | 1624-37 | post-larva | 1189.33 | 825.73 |
| *B. azoricus* | 1624-16 | post-larva | 1200.26 | 794.25 |
| *B. azoricus* | 1624-53 | post-larva | 1589.38 | 951.29 |
| *B. azoricus* | 1626-01 | juvenile | 2170.00 | 1430.00 |
| *B. azoricus* | 1624-15 | juvenile | 2287.17 | 1526.11 |
| *B. azoricus* | 1624-114 | juvenile | 2500.00 | 1600.00 |
| *B. azoricus* | 1629-124 | juvenile | 2600.00 | 1500.00 |
| *B. azoricus* | 1624-115 | juvenile | 2700.00 | 1700.00 |
| *B. azoricus* | 1626-03 | juvenile | 2730.00 | 1120.00 |
| *B. azoricus* | 1626-02 | juvenile | 2860.00 | 1630.00 |
| *B. azoricus* | 1624-112 | juvenile | 2900.00 | 1800.00 |

| | | | | |
|---|---|---|---|---|
| *B. azoricus* | 1624-116 | juvenile | 2900.00 | 1800.00 |
| *B. azoricus* | 1629-123 | juvenile | 3000.00 | 1800.00 |
| *B. azoricus* | 1624-117 | juvenile | 3100.00 | 1900.00 |
| *B. azoricus* | 1624-118 | juvenile | 3100.00 | 2000.00 |
| *B. azoricus* | 1624-113 | juvenile | 3200.00 | 2000.00 |
| *B. azoricus* | 1624-119 | juvenile | 3200.00 | 1800.00 |
| *B. azoricus* | 1624-120 | juvenile | 3200.00 | 2000.00 |
| *B. azoricus* | 1624-125 | juvenile | 3200.00 | 1900.00 |
| *B. azoricus* | 1624-126 | juvenile | 3600.00 | 2100.00 |
| *B. azoricus* | 1626-04 | juvenile | 3630.00 | 2020.00 |
| *B. azoricus* | 1624-121 | adult | 10600.00 | 6000.00 |
| *"B". childressi* | M102 | plantigrade | 383.82 | 406.78 |
| *"B". childressi* | M103 | post-larva | 433.13 | 463.17 |
| *"B". childressi* | M101 | pediveliger | 434.37 | 447.76 |
| *"B". childressi* | M099 | post-larva | 949.75 | 713.60 |
| *"B". childressi* | M098 | post-larva | 1070.29 | 671.88 |
| *"B". childressi* | M100 | post-larva | 1325.54 | 1027.59 |
| *B. puteoserpentis* | 1556-11 | pediveliger / plantigrade | 366.37 | 393.70 |
| *B. puteoserpentis* | 1555-30 | post-larva | 372.42 | 353.10 |
| *B. puteoserpentis* | 1555-27 | plantigrade | 385.47 | 417.85 |
| *B. puteoserpentis* | 1556-36 | post-larva | 391.74 | 306.57 |
| *B. puteoserpentis* | 1555-12 | pediveliger / plantigrade | 392.69 | 424.94 |
| *B. puteoserpentis* | 1556-45 | pediveliger / plantigrade | 396.58 | 362.52 |
| *B. puteoserpentis* | 1555-16 | pediveliger | 399.37 | 420.37 |
| *B. puteoserpentis* | 1555-15 | pediveliger / plantigrade | 403.77 | 417.71 |
| *B. puteoserpentis* | 1555-06 | pediveliger | 404.99 | 427.36 |
| *B. puteoserpentis* | 1556-30 | pediveliger / plantigrade | 405.11 | 433.54 |
| *B. puteoserpentis* | 1555-32 | pediveliger / plantigrade | 405.47 | 410.07 |
| *B. puteoserpentis* | 1556-58 | post-larva | 406.57 | 424.66 |
| *B. puteoserpentis* | 1556-48 | pediveliger / plantigrade | 406.90 | 285.45 |
| *B. puteoserpentis* | 1556-61 | pediveliger | 407.91 | 382.31 |
| *B. puteoserpentis* | 1555-31 | pediveliger | 409.30 | 408.16 |
| *B. puteoserpentis* | 1555-28 | pediveliger | 409.34 | 415.80 |
| *B. puteoserpentis* | 1556-26 | plantigrade | 410.38 | 428.54 |
| *B. puteoserpentis* | 1555-45 | plantigrade | 411.13 | 442.17 |
| *B. puteoserpentis* | 1555-04 | pediveliger | 411.23 | 402.91 |
| *B. puteoserpentis* | 1556-38 | pediveliger / plantigrade | 413.07 | 449.15 |
| *B. puteoserpentis* | 1556-29 | post-larva | 414.18 | 394.26 |
| *B. puteoserpentis* | 1555-14 | post-larva | 414.32 | 394.93 |
| *B. puteoserpentis* | 1556-35 | pediveliger / plantigrade | 414.35 | 426.24 |
| *B. puteoserpentis* | 1555-42 | plantigrade | 417.82 | 413.34 |
| *B. puteoserpentis* | 1555-13 | pediveliger / plantigrade | 418.29 | 413.66 |
| *B. puteoserpentis* | 1556-05 | pediveliger | 418.47 | 463.48 |
| *B. puteoserpentis* | 1556-23 | pediveliger / plantigrade | 420.67 | 374.87 |
| *B. puteoserpentis* | 1556-21 | pediveliger / plantigrade | 420.91 | 402.18 |
| *B. puteoserpentis* | 1556-41 | pediveliger / plantigrade | 421.44 | 420.46 |
| *B. puteoserpentis* | 1556-42 | pediveliger / plantigrade | 427.02 | 426.17 |
| *B. puteoserpentis* | 1556-31 | pediveliger / plantigrade | 429.36 | 458.44 |
| *B. puteoserpentis* | 1556-08 | post-larva | 430.08 | 416.34 |
| *B. puteoserpentis* | 1555-44 | post-larva | 431.15 | 401.41 |
| *B. puteoserpentis* | 1555-03 | post-larva | 432.29 | 447.89 |
| *B. puteoserpentis* | 1556-12 | pediveliger / plantigrade | 432.40 | 435.28 |
| *B. puteoserpentis* | 1556-62 | post-larva | 432.80 | 433.97 |
| *B. puteoserpentis* | 1556-07 | post-larva | 434.02 | 445.35 |
| *B. puteoserpentis* | 1556-43 | pediveliger / plantigrade | 434.19 | 429.49 |
| *B. puteoserpentis* | 1556-32 | pediveliger / plantigrade | 434.36 | 454.08 |
| *B. puteoserpentis* | 1556-28 | pediveliger / plantigrade | 435.47 | 429.78 |

| | | | | |
|---|---|---|---|---|
| *B. puteoserpentis* | 1555-01 | post-larva | 435.53 | 404.12 |
| *B. puteoserpentis* | 1555-05 | plantigrade | 436.44 | 390.65 |
| *B. puteoserpentis* | 1555-02 | post-larva | 439.64 | 493.34 |
| *B. puteoserpentis* | 1556-10 | post-larva | 450.10 | 462.96 |
| *B. puteoserpentis* | 1556-22 | post-larva | 453.32 | 461.34 |
| *B. puteoserpentis* | 1556-40 | post-larva | 455.20 | 402.18 |
| *B. puteoserpentis* | 1556-02 | post-larva | 456.18 | 460.44 |
| *B. puteoserpentis* | 1556-53 | post-larva | 466.25 | 450.42 |
| *B. puteoserpentis* | 1556-06 | post-larva | 470.17 | 445.82 |
| *B. puteoserpentis* | 1556-37 | post-larva | 473.58 | 461.28 |
| *B. puteoserpentis* | 1556-57 | post-larva | 473.80 | 467.54 |
| *B. puteoserpentis* | 1555-18 | post-larva | 476.51 | 440.72 |
| *B. puteoserpentis* | 1556-03 | post-larva | 482.94 | 463.48 |
| *B. puteoserpentis* | 1556-17 | post-larva | 484.55 | 500.04 |
| *B. puteoserpentis* | 1556-44 | post-larva | 501.13 | 453.77 |
| *B. puteoserpentis* | 1556-34 | post-larva | 526.99 | 520.48 |
| *B. puteoserpentis* | 1556-04 | post-larva | 529.17 | 499.98 |
| *B. puteoserpentis* | 1556-33 | post-larva | 541.16 | 475.78 |
| *B. puteoserpentis* | 1556-24 | post-larva | 544.27 | 457.28 |
| *B. puteoserpentis* | 1556-25 | post-larva | 549.36 | 542.46 |
| *B. puteoserpentis* | 1556-09 | post-larva | 557.16 | 524.54 |
| *B. puteoserpentis* | 1556-16 | post-larva | 576.86 | 563.26 |
| *B. puteoserpentis* | 1556-20 | post-larva | 592.62 | 516.92 |
| *B. puteoserpentis* | 1556-15 | post-larva | 594.13 | 448.48 |
| *B. puteoserpentis* | 1555-17 | post-larva | 600.99 | 508.09 |
| *B. puteoserpentis* | 1556-59 | post-larva | 601.97 | 521.19 |
| *B. puteoserpentis* | 1555-25 | post-larva | 602.05 | 466.03 |
| *B. puteoserpentis* | 1555-23 | post-larva | 607.31 | 501.68 |
| *B. puteoserpentis* | 1556-46 | post-larva | 613.12 | 539.58 |
| *B. puteoserpentis* | 1555-33 | post-larva | 618.94 | 509.97 |
| *B. puteoserpentis* | 1555-24 | post-larva | 637.08 | 536.66 |
| *B. puteoserpentis* | 1555-43 | post-larva | 711.47 | 592.00 |
| *B. puteoserpentis* | 1555-10 | post-larva | 716.63 | 586.06 |
| *B. puteoserpentis* | 1556-14 | post-larva | 724.06 | 501.79 |
| *B. puteoserpentis* | 1555-11 | post-larva | 759.12 | 633.82 |
| *B. puteoserpentis* | 1555-35 | post-larva | 823.51 | 546.17 |
| *B. puteoserpentis* | 1556-60 | post-larva | 825.96 | 671.65 |
| *B. puteoserpentis* | 1555-26 | post-larva | 879.14 | 674.50 |
| *B. puteoserpentis* | 1556-19 | post-larva | 886.64 | 718.74 |
| *B. puteoserpentis* | 1555-39 | post-larva | 962.04 | 725.98 |
| *B. puteoserpentis* | 1555-36 | post-larva | 997.00 | 774.75 |
| *B. puteoserpentis* | 1555-38 | post-larva | 998.26 | 841.35 |
| *B. puteoserpentis* | 1556-39 | post-larva | 1092.47 | 788.72 |
| *B. puteoserpentis* | 1555-37 | post-larva | 1126.02 | 889.74 |
| *B. puteoserpentis* | 1556-63 | post-larva | 1178.77 | 656.87 |
| *B. puteoserpentis* | 1555-19 | post-larva | 1192.74 | 910.69 |
| *B. puteoserpentis* | 1556-52 | post-larva | 1212.82 | 878.79 |
| *B. puteoserpentis* | 1555-29 | post-larva | 1221.64 | 938.29 |
| *B. puteoserpentis* | 1555-40 | post-larva | 1284.74 | 943.78 |
| *B. puteoserpentis* | 1556-18 | post-larva | 1325.99 | 903.58 |
| *B. puteoserpentis* | 1556-27 | post-larva | 1384.38 | 905.20 |
| *B. puteoserpentis* | 1556-56 | post-larva | 1510.52 | 718.12 |
| *B. puteoserpentis* | 1556-55 | post-larva | 1803.69 | 1036.59 |
| *B. puteoserpentis* | 1555-20 | post-larva | 1859.55 | 1279.78 |
| *B. puteoserpentis* | 1556-01 | post-larva | 1868.40 | 1277.92 |
| *B. puteoserpentis* | 1555-46 | post-larva | 1952.05 | 1278.18 |
| *B. puteoserpentis* | 1556-54 | post-larva | 1962.02 | 1243.06 |

| | | | | |
|---|---|---|---|---|
| *B. puteoserpentis* | 1556-49 | juvenile | 2053.09 | 1451.25 |
| *B. puteoserpentis* | 1556-51 | juvenile | 2065.05 | 1246.96 |
| *B. puteoserpentis* | 1555-47 | juvenile | 2290.10 | 1508.85 |
| *B. puteoserpentis* | 1542-08 | juvenile | 2451.12 | 1586.87 |
| *B. puteoserpentis* | 1556-50 | juvenile | 2457.04 | 1942.16 |
| *B. puteoserpentis* | 1274-2 | juvenile | 2524.32 | 1395.75 |
| *B. puteoserpentis* | 1542-02 | juvenile | 2690.55 | 1743.98 |
| *B. puteoserpentis* | 1555-48 | juvenile | 2924.96 | 1879.83 |
| *B. puteoserpentis* | 1555-22 | juvenile | 3072.61 | 2060.11 |
| *B. puteoserpentis* | 1542-01 | juvenile | 3264.23 | 2132.32 |
| *B. puteoserpentis* | 1555-21 | juvenile | 3269.13 | 2216.87 |
| *B. puteoserpentis* | 1542-03 | juvenile | 3349.20 | 2254.64 |
| *B. puteoserpentis* | 1542-05 | juvenile | 3367.48 | 2309.16 |
| *B. puteoserpentis* | 1556-64 | juvenile | 3462.94 | 2475.79 |
| *B. puteoserpentis* | 1542-07 | juvenile | 3489.01 | 2288.36 |
| *B. puteoserpentis* | 1542-04 | juvenile | 3665.60 | 2481.33 |
| *B. puteoserpentis* | 1274-1 | juvenile | 3676.82 | 2148.00 |
| *B. puteoserpentis* | 1542-09 | juvenile | 3702.11 | 2467.06 |
| *B. puteoserpentis* | 1542-06 | juvenile | 3735.93 | 2450.20 |
| *B. puteoserpentis* | 1556-47 | juvenile | 3952.60 | 2389.68 |
| *B. puteoserpentis* | 1277-4 | juvenile | 4143.63 | 2375.39 |
| *B. puteoserpentis* | 1277-1 | juvenile | 4510.80 | 2560.52 |
| *B. puteoserpentis* | 1277-3 | juvenile | 4992.68 | 2956.97 |
| *B. puteoserpentis* | 1277-2 | adult | 5162.75 | 2404.32 |
| *B. puteoserpentis* | 812-8 | adult | 5763.25 | 3051.52 |
| *B. puteoserpentis* | 812-7 | adult | 6302.81 | 4049.28 |
| *B. puteoserpentis* | 812-3 | adult | 6818.49 | 4433.88 |
| *B. puteoserpentis* | 812-6 | adult | 7801.08 | 5092.31 |
| *B. puteoserpentis* | 812-2 | adult | 8336.78 | 5192.61 |
| *B. puteoserpentis* | 812-4 | adult | 8893.70 | 5159.90 |
| *B. puteoserpentis* | 812-1 | adult | 10658.47 | 5525.67 |
| *B. puteoserpentis* | 812-5 | adult | 10964.82 | 5942.90 |

**Table S3. FISH probes used in this study.** All probes were labelled with the corresponding fluorophore on the 3' and 5' ends.

| name | target | fluorophore | sequence (5'–3') | reference |
|---|---|---|---|---|
| EUB I | bacteria | 2x Atto 647 or 2x Atto 550 | GCTGCCTCCCGTAGGAGT | [4] |
| EUB II | bacteria | 2x Atto 647 or 2x Atto 550 | GCAGCCACCCGTAGGTGT | [5] |
| EUB III | bacteria | 2x Atto 647 or 2x Atto 550 | GCTGCCACCCGTAGGTGT | [5] |
| Non 338 | bacteria | 2x Atto 550 | ACTCCTACGGGAGGCAGC | [6] |
| BMARt-193 | sulphur-oxidizing bacteria | 2x Atto 550 or 2xAtto 594 | CGAAGGTCCTCCACTTTA | [7] |
| BMARm-845 | methane-oxidizing bacteria | 2x Pacific Blue or 2x Atto 647 | GCTCCGCCACTAAGCCTA | [7] |

**Table S4. Experimental parameters of the SRµCT samples.** cam, camera; eff. pixel, effective pixel size; FOV, field of view.

| sample parameters | | | | | imaging parameters | | | | | | | reconstruction | |
|---|---|---|---|---|---|---|---|---|---|---|---|---|---|
| species | sample identifier | developmental stage | shell length (µm) | shell height (µm) | mag | FOV (mm) | eff. pixel (µm) | cam | # projections | exposure (ms) | energy (keV) | binning | final pixel size |
| *B. puteoserpentis* | 1555-22 | juvenile | 3072.61 | 2060.11 | 20× | 1.8 × 1.8 | 0.65 | CCD | 1200 | 800 | 16 | 2x | 1.3 µm |
| | 1555-47 | juvenile | 2290.09 | 1508.85 | 20× | 1.8 × 1.8 | 0.65 | CCD | 1200 | 800 | 16 | 2x | 1.3 µm |
| | 1555-14 | post-larva | 414.32 | 394.93 | 40× | 0.9 × 0.9 | 0.35 | CCD | 2400 | 2700 | 16 | 2x | 0.7 µm |
| | 1555-30 | post-larva | 372.42 | 353.10 | 40× | 0.9 × 0.9 | 0.35 | CCD | 2400 | 2000 | 16 | 2x | 0.7 µm |
| | 1555-28 | pediveliger | 409.34 | 415.80 | 40× | 0.9 × 0.9 | 0.35 | CCD | 2400 | 2000 | 16 | 2x | 0.7 µm |
| | 1555-27 | pediveliger | 385.47 | 417.85 | 40× | 0.9 × 0.9 | 0.35 | CCD | 2400 | 2000 | 16 | 2x | 0.7 µm |
| *"B". childressi* | M101 | pediveliger | 434.37 | 447.76 | 40× | 0.9 × 0.9 | 0.35 | CCD | 1200 | 1200 | 16 | 2x | 0.7 µm |
| *B. azoricus* | 1643-02 | post-larva | 498.16 | 510.22 | 40× | 0.9 × 0.9 | 0.35 | CCD | 1200 | 1200 | 16 | 2x | 0.7 µm |
| | 1629-124 | juvenile | 2600 | 1500 | 20× | 1.8 × 1.8 | 0.65 | CCD | 1200 | 1500 | 26,1 | 2x | 1.3 µm |
| | 1624-121 | adult | 10600 | 6000 | 5× | 6.6 × 4.9 | 1.3 | cmos | 2400 | 50 | 25,9 | 4x | 5.1 µm |
| | 1626-01 | juvenile | 2170 | 1430 | 10× | 3.6 x 3.6 | 1.2 | CCD | 1200 | 570 | 25 | 2x | 2.4 µm |

**Table S5. Shell size comparisons of the three *Bathymodiolus* species. *"B". childressi* pediveliger / plantigrade individuals were the smallest and *B. azoricus* the largest.** sd, standard deviation.

| | ***B. azoricus*** | | | | | |
|---|---|---|---|---|---|---|
| | shell length | | | shell height | | |
| | mean (µm) | sd | n | mean (µm) | sd | ***n*** |
| pediveliger / plantigrade | 490.47 | 39.15 | 31 | 527.89 | 22.36 | 31 |
| post-larvae | 649.98 | 194.79 | 67 | 612.74 | 103.72 | 67 |
| juvenile | 2735.43 | 578.02 | 5 | 1545.22 | 326.75 | 5 |
| | ***"B". childressi*** | | | | | |
| | shell length | | | shell height | | |
| | mean (µm) | sd | n | mean (µm) | sd | ***n*** |
| pediveliger / plantigrade | 409.09 | 35.74 | 2 | 427.27 | 28.98 | 2 |
| post-larvae | 944.67 | 375.29 | 4 | 719.06 | 233.04 | 4 |
| juvenile | / | / | / | / | / | / |
| | ***B. puteoserpentis*** | | | | | |
| | shell length | | | shell height | | |
| | mean (µm) | sd | n | mean (µm) | sd | n |
| pediveliger / plantigrade | 412.38 | 17.28 | 32 | 413.50 | 34.73 | 32 |
| post-larvae | 782.08 | 431.85 | 65 | 616.36 | 238.85 | 65 |
| juvenile | 4556.66 | 2430.18 | 32 | 2786.70 | 1316.99 | 32 |

**Table S6. Relative organ sizes in *B. puteoserpentis.* The gills grow substantially over the course of development and the digestive system shrinks between the post-larva and juvenile stages.**

| tissue | pediveliger relative volume (%) | plantigrade relative volume (%) | post-larva relative volume (%) | juvenile / adult relative volume (%) |
|---|---|---|---|---|
| soft body | 100 | 100 | 100 | 100 |
| gills | 4.1 | 23.0 | 23.1 | 56.8 |
| foot | 11.5 | 20.4 | 25.4 | 16.3 |
| central nerve system | 4.1 | 9.0 | 7.3 | 1.2 |
| lipid vesicles | 12.8 | 3.9 | 1.5 | absent |
| digestive system | 29.4 | 37.8 | 35.4 | 9.0 |
| stomach and digestive gland | 1.6 | 5.0 | 4.0 | 7.2 |
| velum | 32.2 | absent | absent | absent |
| adductor muscle | 15.0 | 7.5 | 6.6 | 6.1 |
| retractor muscle | 3.7 | 2.3 | 2.2 | 3.3 |

**Table S7. Bacterial densities are highest in adult mussels. Host and bacterial cell areas were calculated from TEM images and set into relation to estimate bacterial densities.**

| developmental stage | averaged symbiont area per host cell (%) | minimum symbiont area per host cell (%) | maximum symbiont area per host cell (%) | standard deviation | number of host cells / number of mussels |
|---|---|---|---|---|---|
| pediveliger | absent | absent | absent | absent | absent |
| plantigrade | 13.7 | 2.0 | 23.2 | 6.3 | 12 / 2 |
| post-larva | 19.4 | 12.2 | 29.0 | 5.0 | 10 / 2 |
| adults | 24.4 | 15.6 | 32.1 | 6.1 | 6 / 2 |

**Table S8. Degree of digestive system reduction in bivalves living in chemosynthetic systems.** =, normally developed; -, reduced morphology; /, total disappearance; n.d., no data.

| | ***B. azoricus* [6]** | ***B. brooksi* [7]** | ***B. heckerae* [7]** | ***B. thermophilus* [8]** | ***Solemya reidi* [9]** | ***Calyptogena magnifica* [10]** |
|---|---|---|---|---|---|---|
| labial palps | = | = | - | - | - | - |
| intestine | - | - | - | - | / | - |
| stomach | - | - | - | - | / | - |
| crystalline style and style sac | = | n.d. | n.d. | - | / | - |
| digestive tubule | - | - | - | - | / | - |
| digestive gland secretory cells | = | n.d. | n.d. | = | / | - |
| digestive system | - | - | - | - | / | - |

### Table S9 FISH hybridisation buffer

| | **30% formamide (ml)** | **35% formamide (ml)** |
|---|---|---|
| 5 M NaCl | 1.08 | 1.08 |
| 1 M TrisHCl | 0.12 | 0.12 |
| Formamide | 1.80 | 2.10 |
| H2O | 3.00 | 2.70 |
| 20% SDS | 0.003 | 0.003 |

### Table S10 FISH washing buffer

| | **30% formamide (ml)** | **35% formamide (ml)** |
|---|---|---|
| 5 M NaCl | 1.02 | 0.70 |
| 1 M TrisHCl | 1.00 | 1.00 |
| 0.5 M EDTA | 0.50 | 0.50 |
| H2O | add to 50 ml | add to 50 ml |
| 20% SDS | 0.025 | 0.025 |

### Supplementary figures

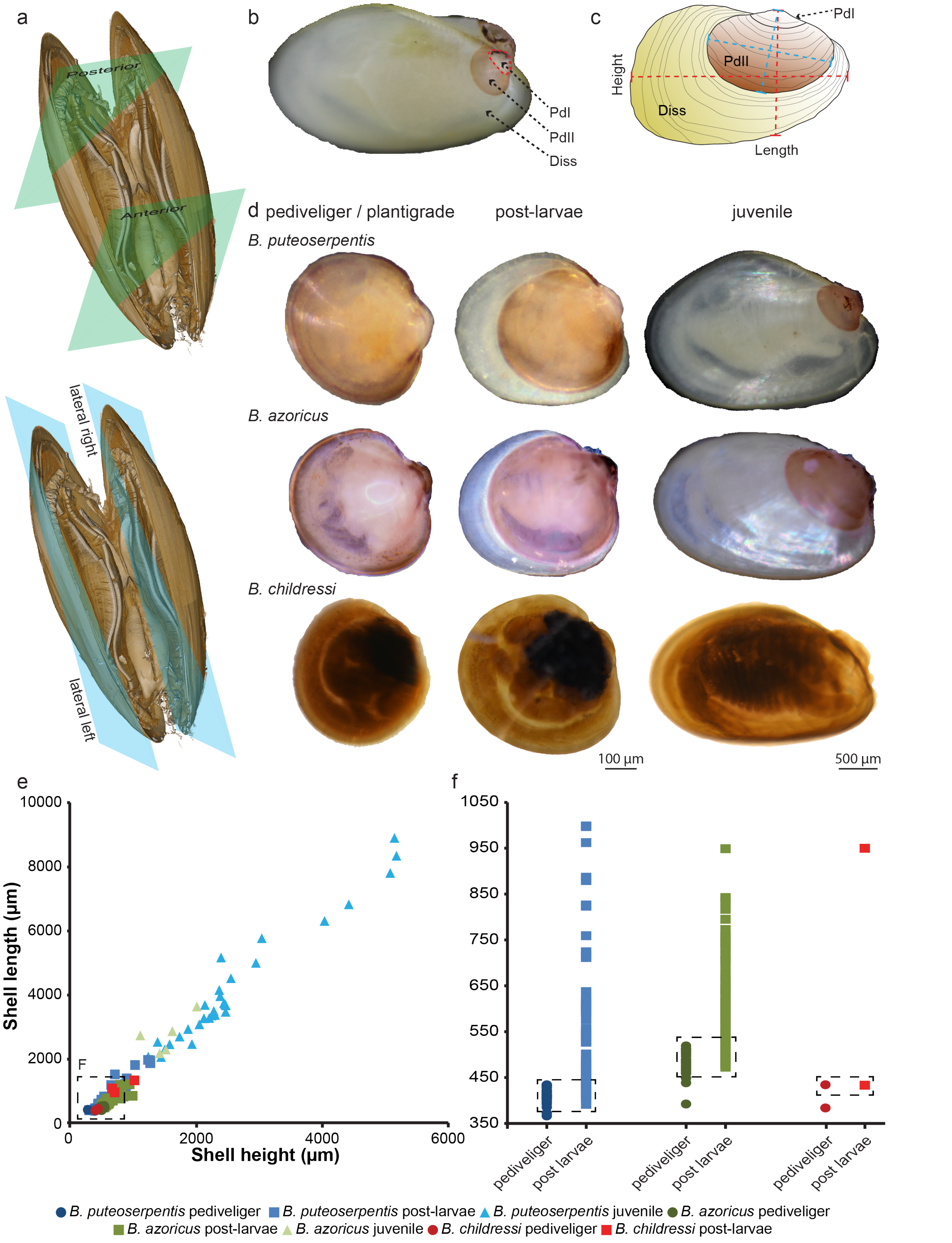

**Figure S1. Sample overview and shell dimensions.**
a) Volume rendering of a *B. azoricus* mussel of 3 cm length showing representations of two different directions of sectioning (anterior to posterior and lateral left to lateral right). b) Visualization of the three different shell types of *Bathymodiolus* mussels. c) Schematic drawing of the different shell types and the directions of shell measurement. d) Examples of the three analysed *Bathymodiolus* species at three developmental stages. e) Plot of shell length vs. shell height, showing overlap between different developmental stages. f) Distribution of shell length for pediveligers and post-larvae. The dashed boxes show the overlap between pediveliger and post-larva. PdI, prodissoconch I; PdII, prodissoconch II; dis, dissoconch.

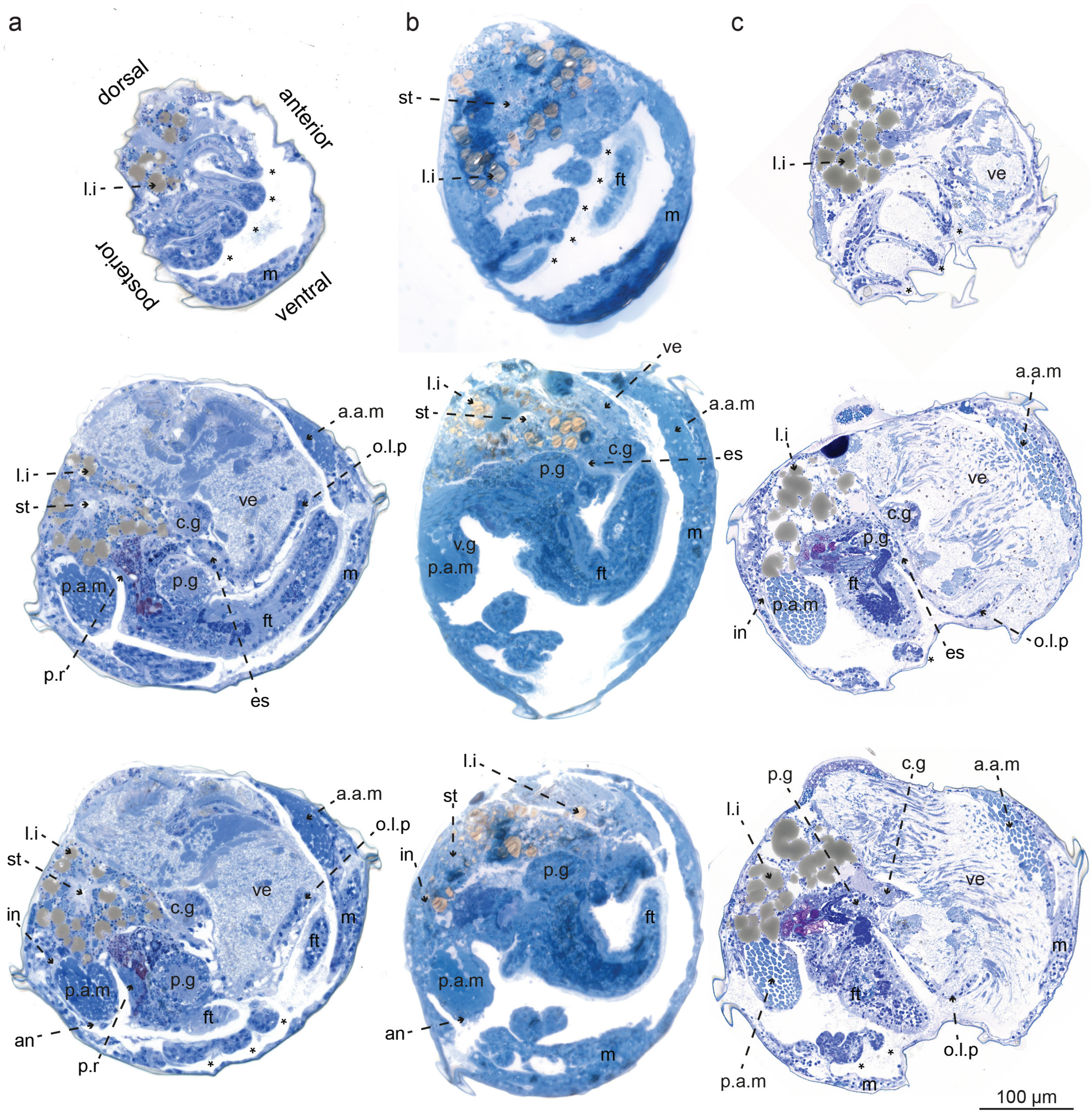

**Figure S2. Morphological comparison of *B. puteoserpentis* (a), *"B". childressi* (b) and *B. azoricus* (c) pediveliger.**
From top to bottom, representative micrographs at incremental locations along the lateral–lateral (left–right) axis presenting the detailed morphology of the three analysed species. All major organs, including the foot, gills, velum, digestive system and central nervous system, are similarly developed in the three species. a.a.m, anterior adductor muscle; c.g, cerebral ganglion; es, esophagus; ft, foot; in, intestine; l.i, lipid vesicles; l.l.p, lower labial palp; m, mantle; p.a.m, posterior adductor muscle; p.g, pedal ganglia; r.m, retractor muscle; s.s, style sac; st, stomach; u.l.p, upper labial palp; ve, velum; *, gill filament.

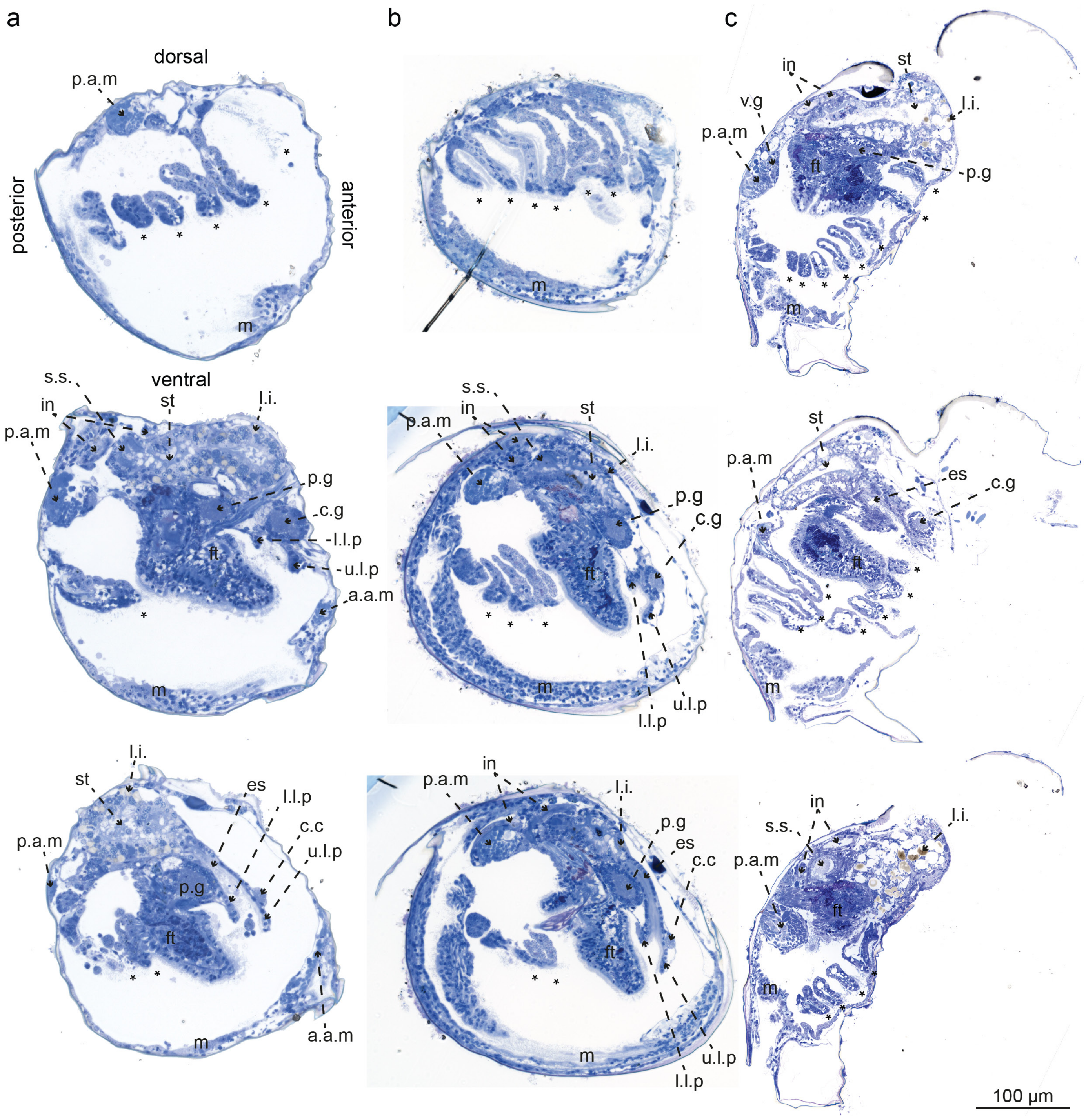

**Figure S3. Morphological comparison of *B. puteoserpentis* (a), *"B". childressi* (b) and *B. azoricus* (c) post-larva.**
From top to bottom, representative micrographs at incremental locations along the lateral–lateral (left–right) axis in post-larvae presenting the detailed morphology of the three analysed species. Only the number of gill filaments differs between the three species. a.a.m, anterior adductor muscle; c.c, cerebral commissure; c.g, cerebral ganglion; es, esophagus; ft, foot; g.s, gastric shield; in, intestine; l.i, lipid inclusion; l.l.p, lower labial palp; m, mantle; p.a.m, posterior adductor muscle; p.g, pedal ganglia; r.m, retractor muscle; s.s, style sac; st, stomach; u.l.p, upper labial palp; v.g, visceral ganglion and *, gill filament.

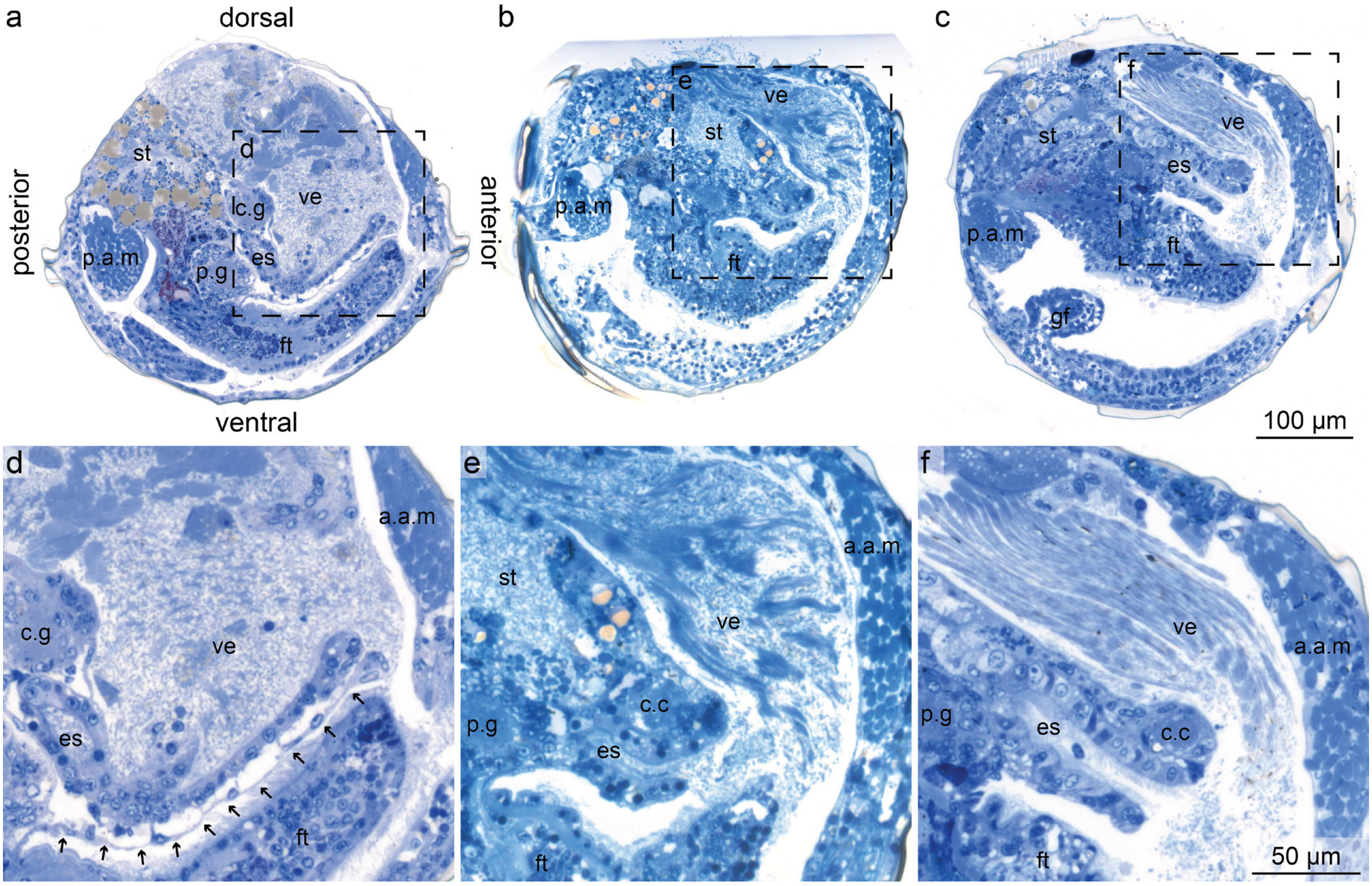

**Figure S4. As metamorphosis in *B. puteoserpentis* progresses, velum degradation begins at the most posterior part of the velum.**
In a first step, the velum membrane (indicated by black arrows in d) is degraded, followed by the degradation of the velar tissue (e,f). Dashed boxes in a–c are shown magnified in d–f. The same morphological pattern was also observed in *B. azoricus* and *"B". childressi.* a.a.m, anterior adductor muscle; c.c, cerebral commissure; c.g, cerebral ganglion; es, esophagus; ft, foot; p.a.m, posterior adductor muscle; p.g, pedal ganglia; st, stomach; ve, velum and *, gill filament.

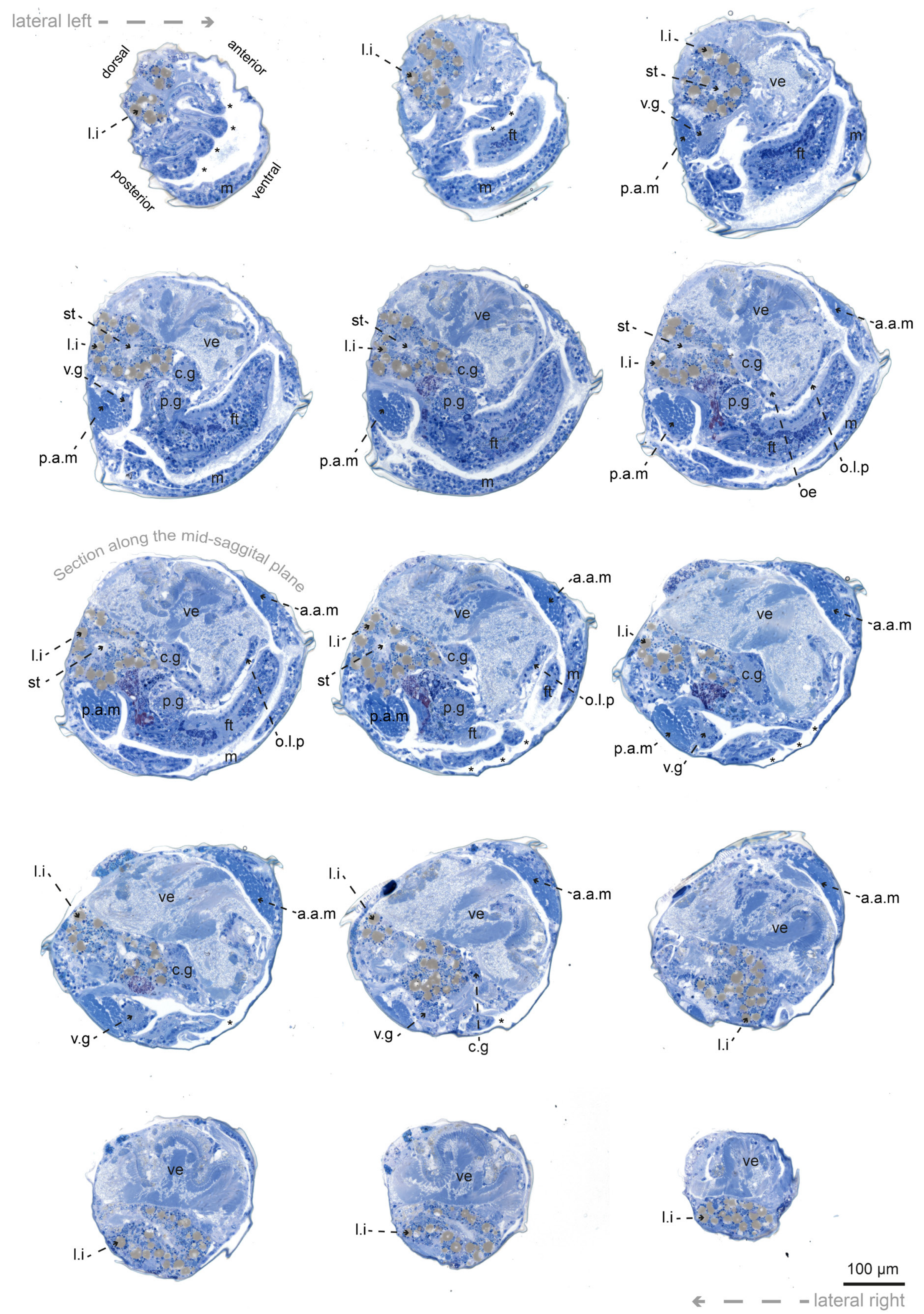

**Figure S5. Serial sections of a *B. puteoserpentis* pediveliger.**
From top left, to bottom right, sections stained with toluidine blue at incremental locations along the sagittal axis. a.a.m, anterior adductor muscle; c.g. cerebral ganglion: ft. foot; l.i, lipid inclusion: m, mantle; o.l.p, oral labial palp; p.a.m, posterior adductor muscle; p.g, pedal ganglia; st, stomach; ve, velum; v.g, visceral ganglion and *, gill filament.

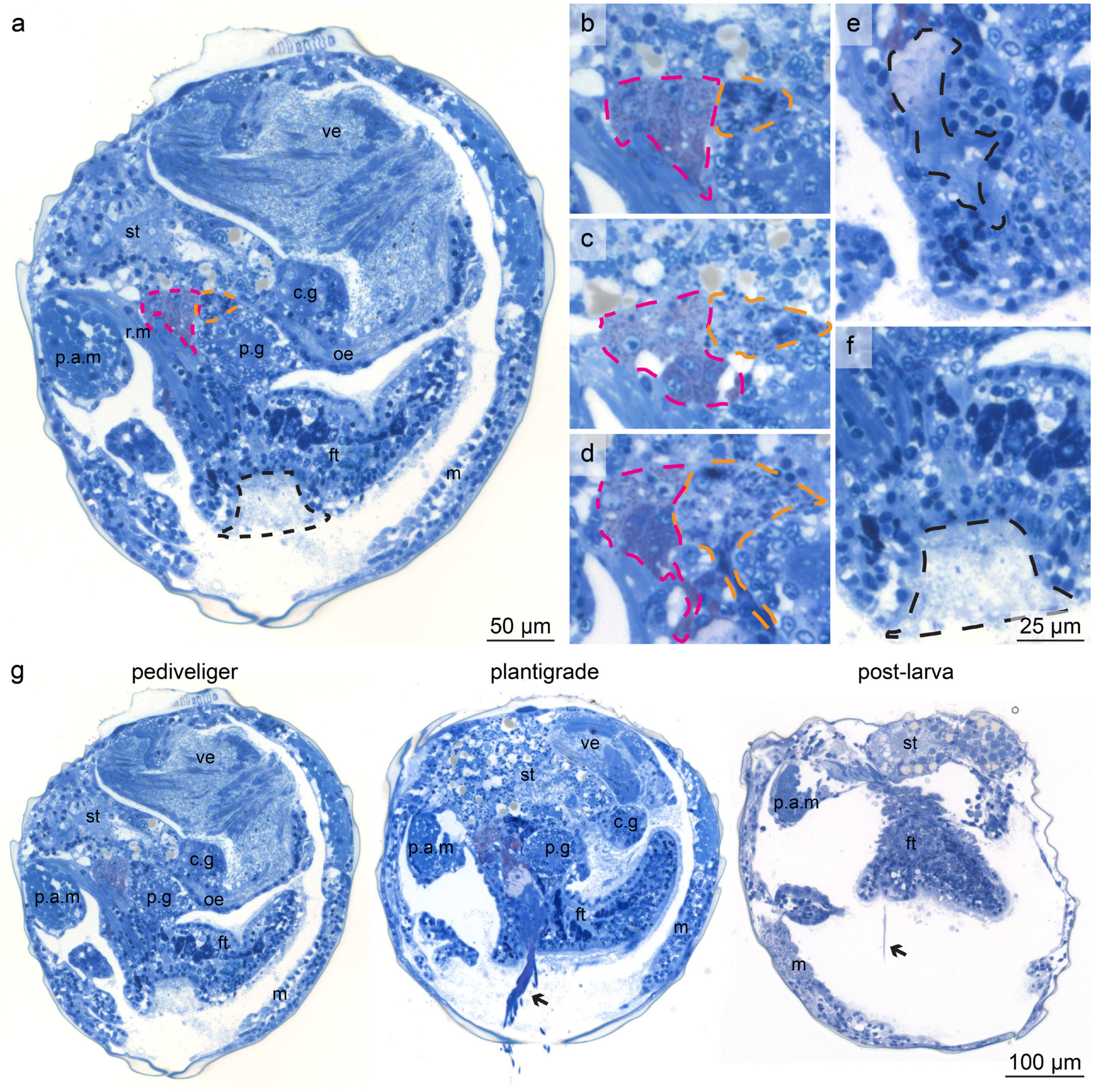

**Figure S6. The *B. puteoserpentis* foot is well developed already during the pediveliger stage.**
In the pediveliger stage, three glands can be identified (a-f): the purple gland (outlined in magenta), the white gland (outlined in orange) and the byssus gland and duct (outlined in black). During metamorphosis, the byssus threads (indicated by black arrows; g) were produced. c.g, cerebral ganglion; ft, foot; m, mantle; oe, oesophagus; p.a.m, posterior adductor muscle; p.g, pedal ganglion; r.m, retractor muscle; st, stomach and ve, velum.

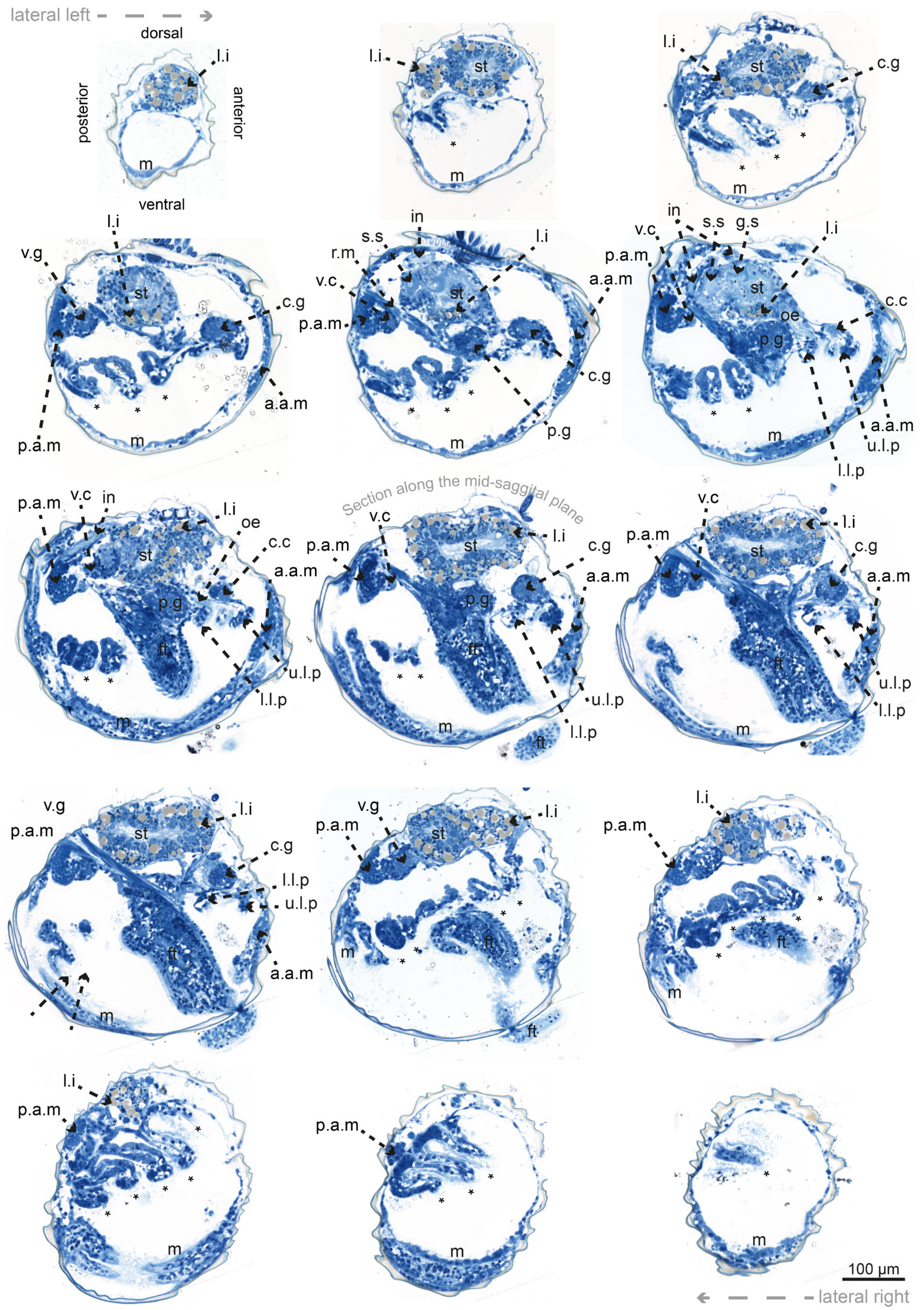

**Figure S7. Serial sections of a *B. puteoserpentis* plantigrade.**
From top left, to bottom right, micrographs at incremental locations along the sagittal axis. a.a.m, anterior adductor muscle; c.g, cerebral ganglion; es, esophagus; ft, foot; g.s, gastric shield; in, intestine; l.i, lipid inclusion; l.l.p, lower labial palp; m, mantle; p.a.m, posterior adductor muscle; p.g, pedal ganglia; r.m, retractor muscle; s.s, style sac; st, stomach; u.l.p, upper labial palp; ve, velum; v.g, visceral ganglion and *, gill filament.

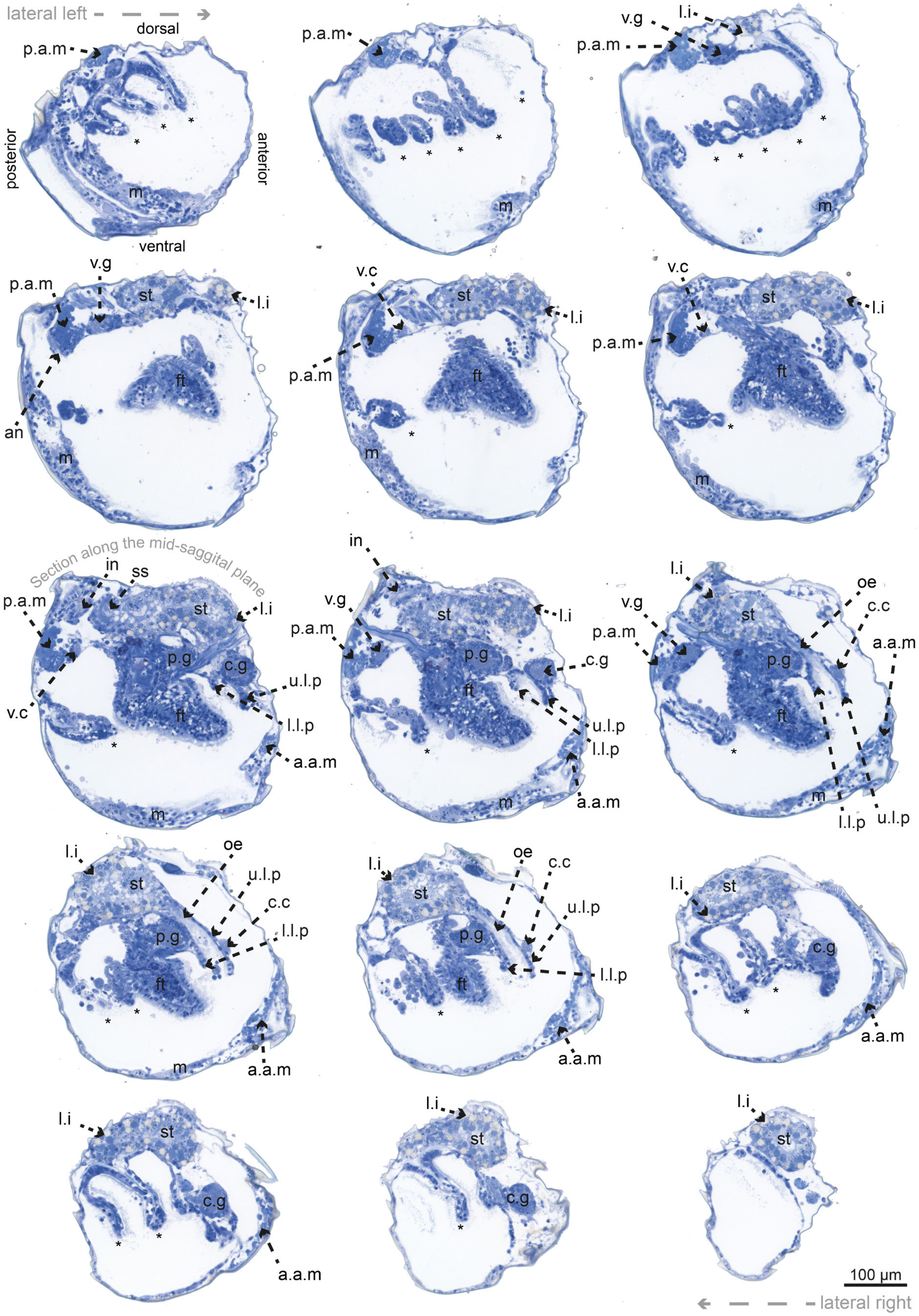

**Figure S8. Serial sections of a *B. puteoserpentis* post-larva.**
From top left, to bottom right, sections stained with toluidine blue at incremental locations along the sagittal axis. a.a.m, anterior adductor muscle; c.g, cerebral ganglion; es, esophagus; ft, foot; g.s, gastric shield; in, intestine; l.i, lipid inclusion; l.l.p, lower labial palp; m, mantle; p.a.m, posterior adductor muscle; p.g, pedal ganglia; r.m, retractor muscle; s.s, style sac; st, stomach; u.l.p, upper labial palp; ve, velum; v.g, visceral ganglion and *, gill filament.

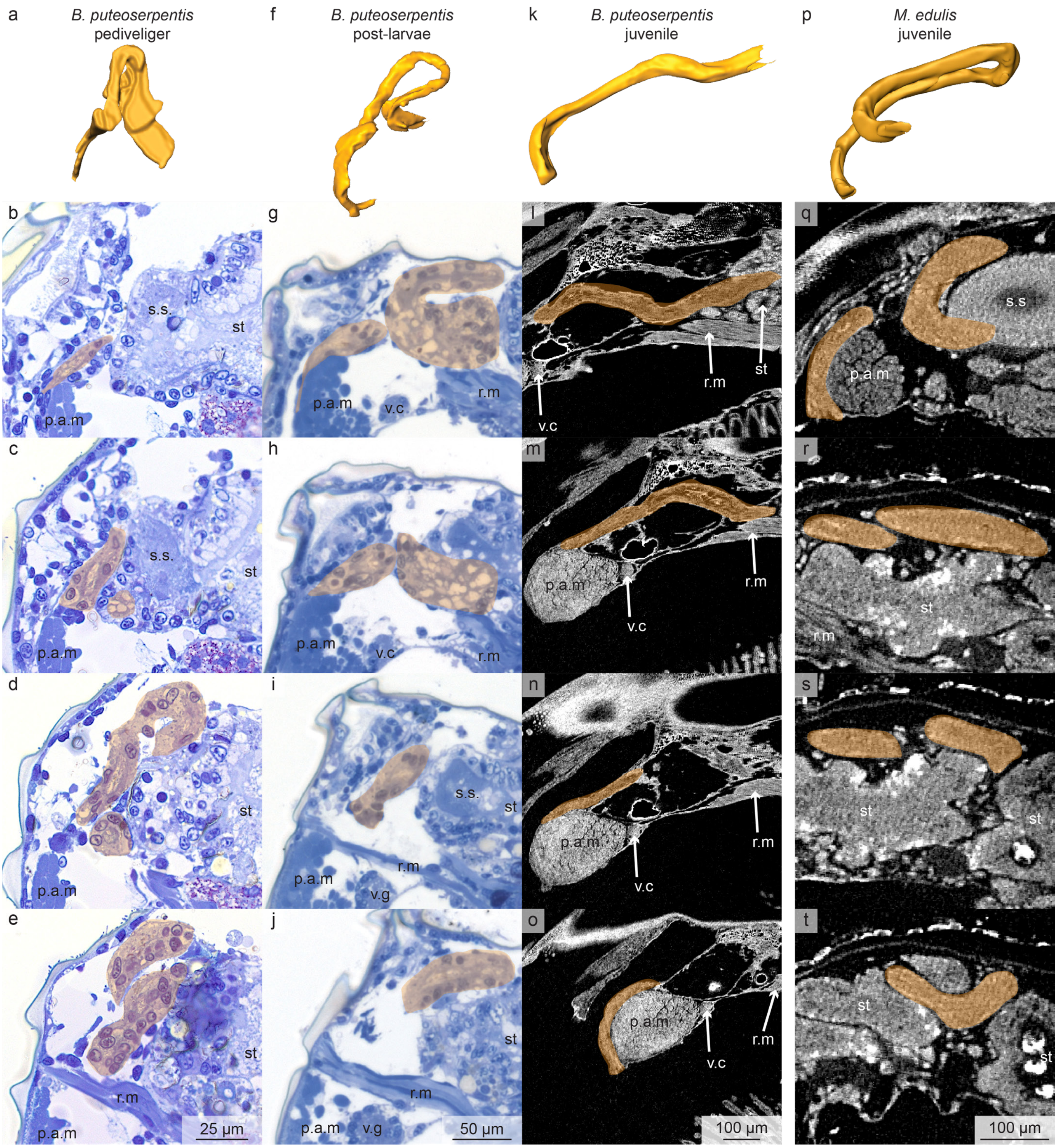

**Figure S9. The looped intestine of the *B. puteoserpentis* pediveliger straightens after metamorphosis in comparison to *M. edulis* were it stays looped.** 3D reconstructions of the intestine (a, f and k) show the morphological change over the course of development in *B. puteoserpentis*. Representative serial sections (top to bottom) show a looped intestine in the pediveliger (b–e) and post-larva (g–j). The straight intestine is shown by visual slices through SRµCT data from a *B. puteoserpentis* juvenile (l–o). In contrast the intestine of a juvenile *M. edulis* mussel stays looped (p, q–t). The intestine is indicated by orange coloration in the section series and SRµCT data. Note the different scales for the pediveliger, post-larva and juvenile mussel. p.a.m, posterior adductor muscle; r.m, retractor muscle; s.s, style sac; st, stomach; v.c, visceral commissure and v.g, visceral ganglion.

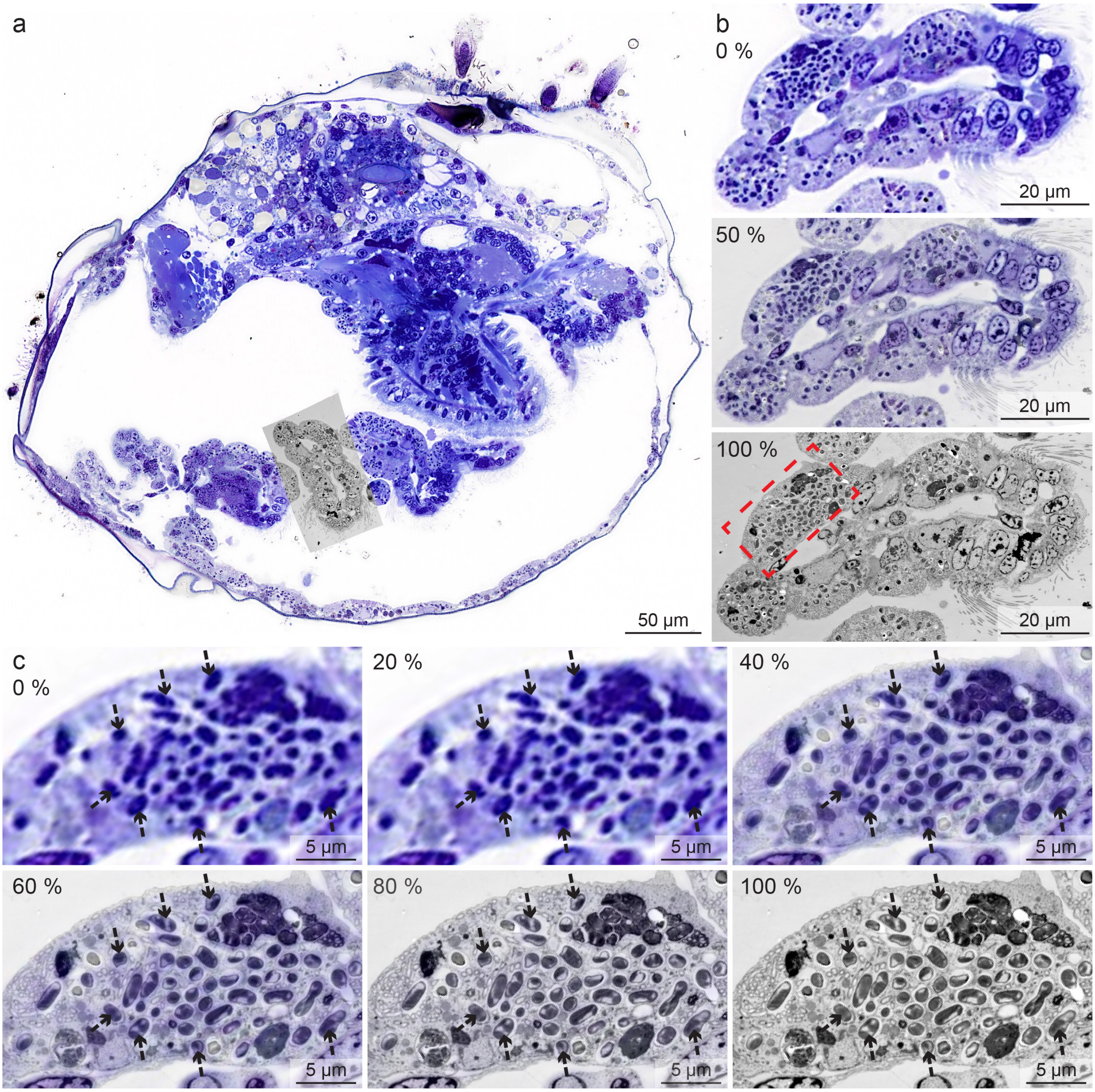

**Figure S10. Light microscopic analysis predicts the presence of symbionts, and can be verified with correlative TEM.**
The overview image (a) shows a cross section of a *B. puteoserpentis* plantigrade which is already colonized by symbionts. Overlaying LM and TEM data of the same section and area allows identification of MOX symbionts as dark blue dots in the LM data (b,c). The percentages are showing value of transparency of the TEM-image. The LM and TEM images were co-registered by using morphological landmarks.

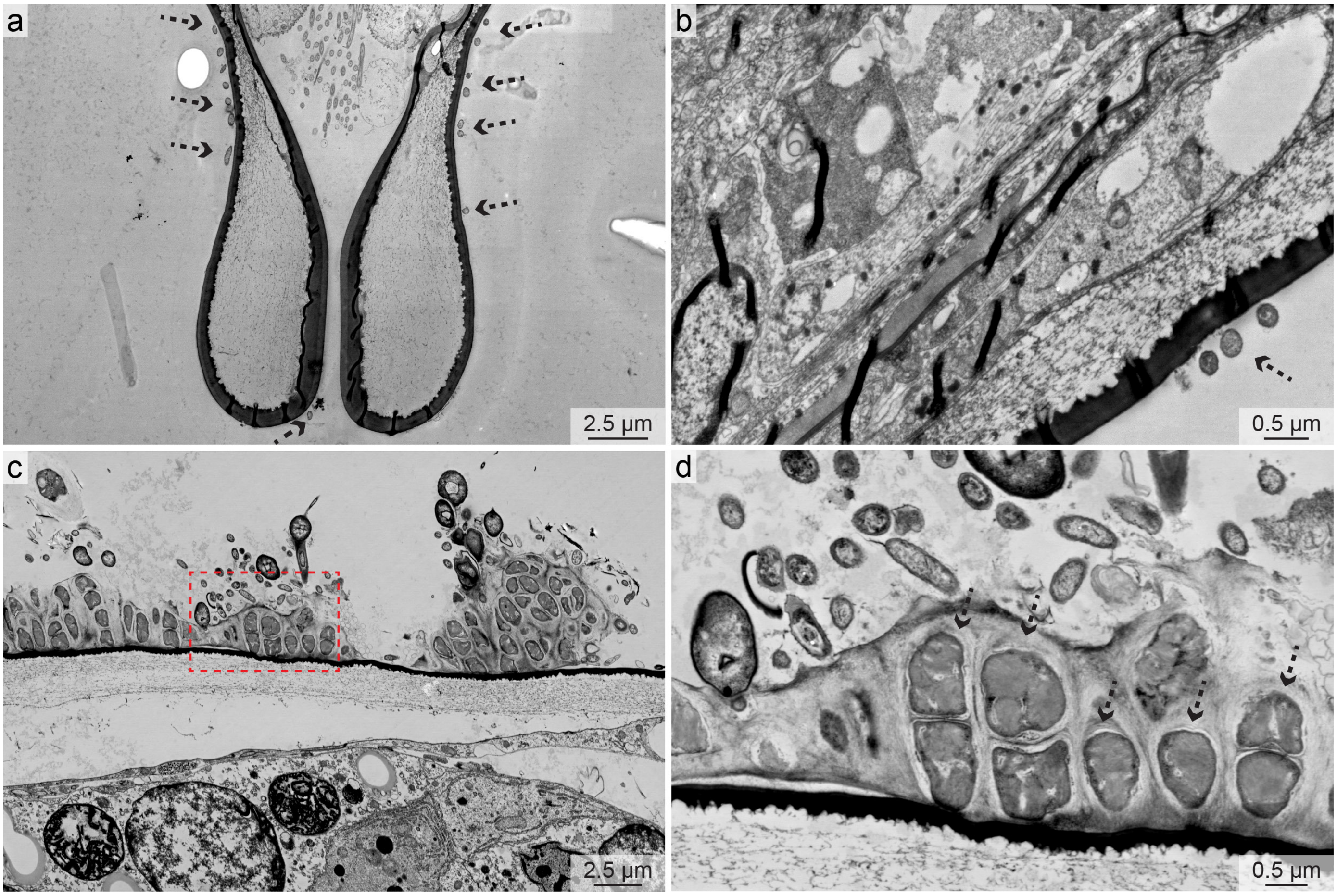

**Figure S11. SOX- and MOX-like bacterial morphotypes were identified on the shells of B. puteoserpentis pediveliger (a and b) and post-larvae (c and d).** TEM data of aposymbiotic pediveliger shows bacteria on the shell that have a similar morphotype to the SOX and MOX symbionts which are indicated byv black dashed arrows.

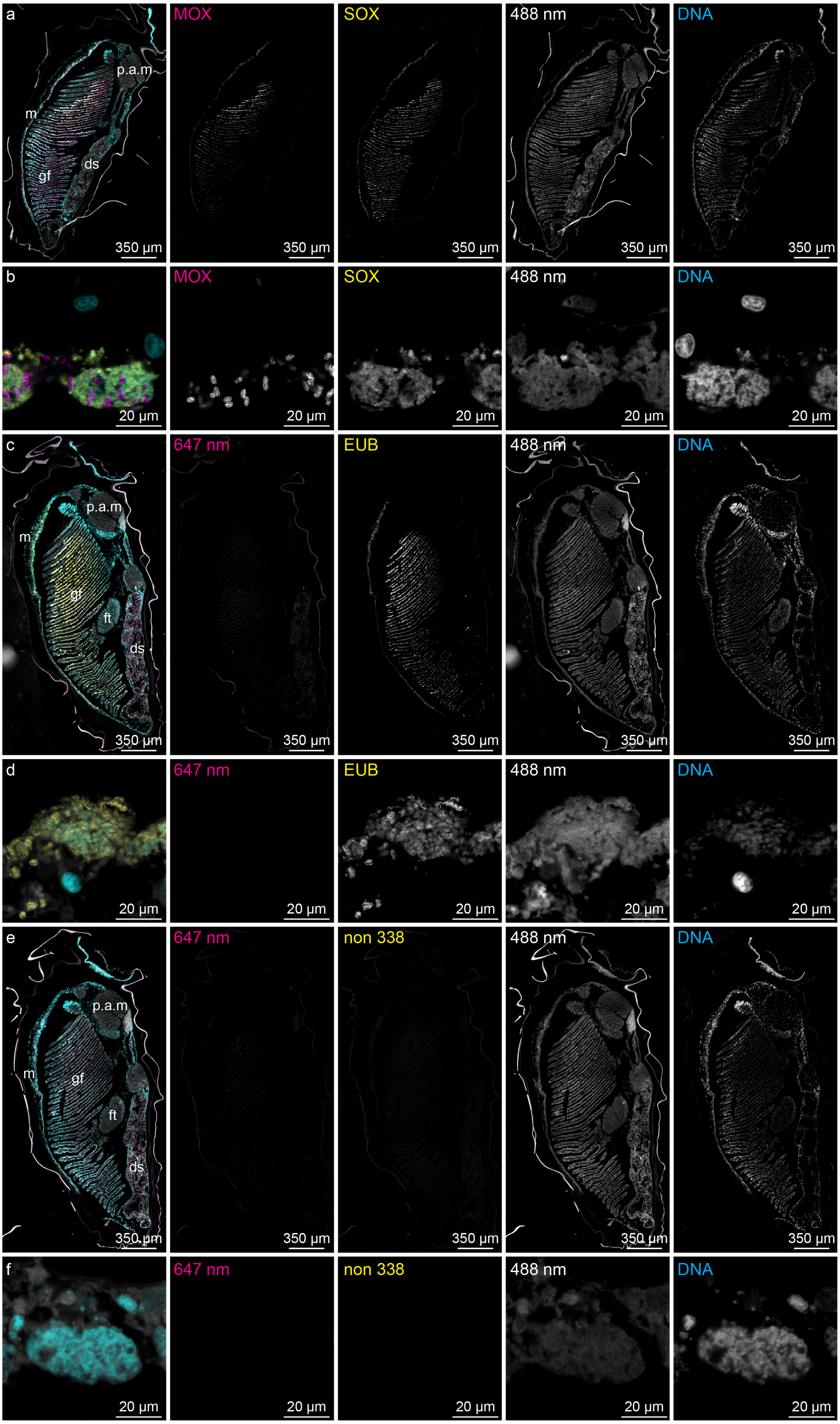

**Figure S12. Fluorescence *in situ* hybridization (FISH) of specific symbionts and controls in cross sections of a** *B. puteoserpentis* **juvenile mussel.** Specific oligonucleotides targeting the rRNA of SOX and MOX bacteria show symbiont distributions in the gills and the mantle (a). SOX symbionts are represented in yellow and MOX symbionts in magenta. The 488-nm and 647-nm channels were used to detect autofluorescence of the host tissue. DAPI was used as a DNA counterstain (represented in cyan). Both symbiont types colonize the same bacteriocytes (b; magnified region of a). As a positive control, the EUB I-III probes which stain all eubacteria were used (c and d; represented in yellow). The EUB I-III signal shows the same symbiont distribution as seen in a and b. As a negative control, the NON 338 probe was used (e and f; represented in yellow), which shows no signal. ds, digestive system; ft, foot: gf, gill filament; m, mantle and p.a.m, posterior adductor muscle

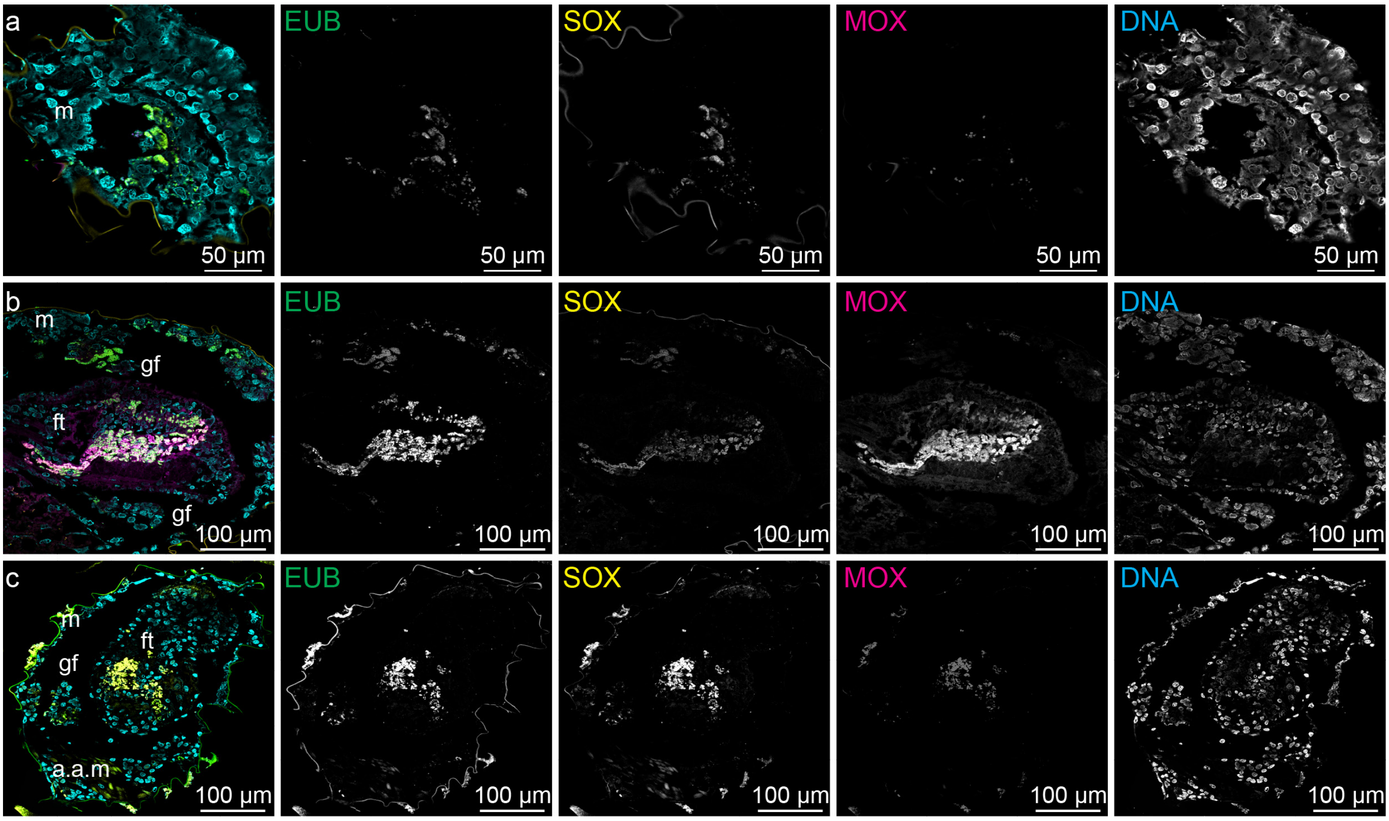

**Figure S13. Fluorescence *in situ* hybridization (FISH) of symbionts in cross sections of *B. puteoserpentis* plantigrades and post-larvae.**
Specific oligonucleotides targeting the rRNA of SOX and MOX bacteria show symbiont distributions in the mantle, foot and gills of a *B. puteoserpentis* post-larva (**a** and **b**). The SOX are represented in yellow and the MOX in magenta. As a positive control, we used the EUB I-III probes (green) to target all bacteria. As a DNA counterstain, DAPI was used (represented in cyan). Both SOX and MOX bacteria were present in all epithelial tissues. The gill filaments of a *B. puteoserpentis* plantigrade (**c**) is only colonized by SOX. a.a.m, anterior adductor muscle; ft, foot; gf, gill filament and m, mantle

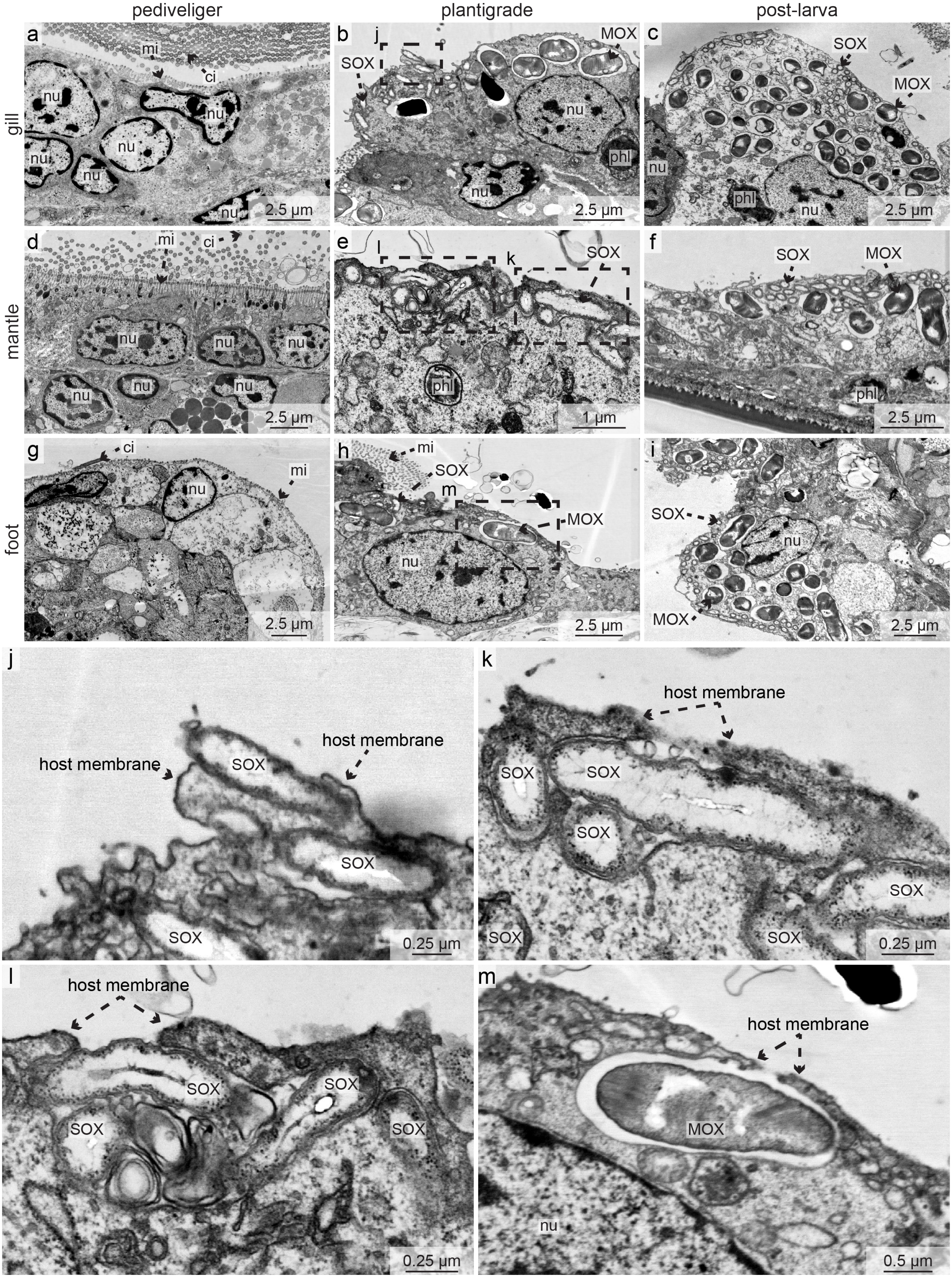

**Figure S14. Symbiont colonization starts during the plantigrade stage in *Bathymodiolus puteoserpentis*.**
TEM micrographs of different epithelial tissues and their state of symbiont colonization: gills (a–c), mantle (d–f) and foot (g–i) of individuals at the pediveliger (a, d and g), plantigrade (b, e and h) and post-larva stage (c, f and i). All epithelial tissues of the pediveliger stage are aposymbiotic (a, d and g). In the plantigrade stage, the colonization of both symbiont types is still ongoing (b, e and h). In the post-larva stage, all epithelial tissues are colonized by both symbiont types (c, f and i). Dashed boxes indicate regions in which symbionts are actively colonizing epithelial tissue (shown magnified in j – m). ci, cilia; mi, microvilli; MOX, methane-oxidizing symbiont; nu, nucleus; phl, phagolysosome; SOX, sulphur-oxidizing symbiont.
